# Quantitation of single action potential-evoked Ca^2+^ signals in CA1 pyramidal neuron presynaptic terminals

**DOI:** 10.1101/337816

**Authors:** Emily Church, Edaeni Hamid, Simon Alford

## Abstract

Presynaptic Ca^2+^ evokes exocytosis, endocytosis, and short-term synaptic plasticity. However, Ca^2+^ flux and interactions at presynaptic molecular targets are difficult to determine, because imaging has limited resolution. We measured single varicosity presynaptic Ca^2+^ using Ca^2+^ dyes as buffers, and constructed models of Ca^2+^ dispersal. Action potentials evoked Ca^2+^ transients (peak amplitude, 789±39 nM, within 2 ms of stimulation; decay times, 119±10 ms) with little variation when measured with low-affinity dye. Endogenous Ca^2+^ buffering capacities, action potential-evoked free [Ca^2+^]¡ and total amounts entering terminals were determined using high-affinity Ca^2+^ dyes to buffer Ca^2+^ transients. These data constrained Monte Carlo (MCell) simulations of Ca^2+^ entry, buffering, and removal. Data were well-fit with simulations of experimentally-determined Ca^2+^ fluxes, buffered by simulated Calbindin_28K_. Simulations were consistent with clustered Ca^2+^ entry followed within 2 ms by diffusion throughout the varicosity. Repetitive stimulation caused free varicosity Ca^2+^ to sum. However, simulated in nanometer domains, its removal by pumps and buffering was negligible, while diffusion rates were high. Thus, Ca^2+^ within tens of nanometers of entry, did not accumulate during sequential stimuli. A model of synaptotagmin1-Ca^2+^ binding indicates that even with 10 μM free varicosity Ca^2+^, synaptogmin1 must be within tens of nanometers of channels to ensure occupation of all its Ca^2+^ binding sites. Repetitive stimulation, which evokes short-term synaptic enhancement, does not modify probabilities of Ca^2+^ fully occupying synaptotagmin1’s C2 domains, suggesting that enhancement is not mediated by Ca^2+^-synaptotagmin1. We conclude that at spatio-temporal scale of fusion machines, Ca^2+^ necessary for their activation is diffusion dominated.

## Introduction

Presynaptic [Ca^2+^] entry through voltage-gated Ca^2+^ channels (VGCCs) causes exocytosis (Katz and Miledi, 1967). Exocytosis may require just one VGCC (Stanley, 1993; Bucurenciu et al., 2010; Weber et al., 2010; Eggermann et al., 2012; Scimemi and Diamond, 2012), or their clustering at microdomains (Llinas et al., 1992; Shahrezaei and Delaney, 2004; Oheim et al., 2006). Additionally, Ca^2+^ plays other roles in presynaptic terminals including modifying repetitive neurotransmission (Zucker, 1989; Zucker and Regehr, 2002) and receptor-mediated neuromodulation (Yoon et al., 2007; Gerachshenko et al., 2009).

Presynaptic Ca^2+^ transients are resolvable with Ca^2+^ dyes (DiGregorio and Vergara, 1997; Cochilla and Alford, 1998; Koester and Sakmann, 2000). However, Ca^2+^ is utilized too locally and rapidly to be imaged within the dimensions of Ca^2+^-binding molecules that cause exocytosis - (Adler et al., 1991; Sabatini and Regehr, 1996). Nevertheless, dyes can quantify presynaptic Ca^2+^ entry, buffering and removal (Neher and Augustine, 1992; Koester and Sakmann, 2000; Jackson and Redman, 2003; Brenowitz and Regehr, 2007). This approach requires calibration of dye concentrations and Ca^2+^ binding properties within cells.

CA1 pyramidal neuron somata and dendrites contain approximately 40 *μ*M of the Ca^2+^ binding protein calbindin_28K_ (Baimbridge et al., 1992; Müller et al., 2005) which may represent their dominant buffer (Yi, 2013) – although other buffers will have an impact. Indeed, calmodulin is present at presynaptic terminals (Llinas et al., 1991; Hinds et al., 2003). Buffering characteristics of these EF hand proteins have been characterized *in vitro* (Nägerl et al., 2000; Faas and Mody, 2012) and by modeling *in situ* (Schmidt et al., 2005), allowing their impact on local Ca^2+^ signaling to be modeled.

Synaptotagmin1 is idely considered the principal Ca^2+^ sensor for evoked release in pyramidal synapses (Geppert et al., 1994). In many synapses, exocytosis requires close association (<100 nm) between Ca^2+^ entry and synaptotagmin (Adler et al., 1991; Martens et al., 2007; Chapman, 2008; Südhof and Rizo, 2011). Ca^2+^ buffers modify Ca^2+^ diffusion, and Ca^2+−^ synaptotagmin interactions. Though widely accepted (DiGregorio and Vergara, 1997; Koester and Sakmann, 2000) this is poorly characterized in presynaptic terminals. Synaptotagmin 1 (syt1) has two Ca^2+^ binding domains (C2A and C2B) (Perin et al., 1991) which bind 3 and 2 Ca^2+^ ions and is the Ca^2+^ sensor in CA1 axons. However, it is unclear whether all syt1 Ca^2+^ sites must bind Ca^2+^ given very low affinities for the C2B domain. Indeed, only one domain is necessary for membrane interaction (Davletov and Sudhof, 1993), although this low affinity increases on association with lipid membranes containing phosphatidylinositol 4,5-bisphosphate PI(4,5)P2 (Radhakrishnan et al., 2009; van den Bogaart et al., 2012). To understand how presynaptic Ca^2+^ interacts with synaptotagmin, we have quantified [Ca^2+^]_i_ in CA1 presynaptic terminals during single action potentials. This allowed us to simulate action-potential-evoked Ca^2+^ entry, binding, buffering and dispersal at individual terminals using Monte Carlo simulation (MCell (Kerr et al., 2008) and investigate its interaction with syt1 at resolutions that evoke exocytosis.

## Results

### Loading cells and resting Ca^2+^ concentrations in axonal varicosities

To measure presynaptic Ca^2+^, dye was introduced to CA1 pyramidal neurons from somatic whole cell pipettes containing Ca^2+^-sensitive dye (Fig 1B), and Alexa 594 hydrazide (250 *μ*M; Fig 1A). After 20 mins, the axon was traced by imaging Alexa 594 (Hamid et al., 2014). We initially used Fluo-4 (1mM) as a Ca^2+^ sensor. Both dyes diffused into axon varicosities at similar rates (Fig 1C) and Alexa 594 fluorescence was used to measure dye concentration. We assume co-diffusion of the dyes which have similar molecular weights (736 and 737 g·mol^−1^). Thus, dye concentrations were calculated throughout each experiment (methods). MCell simulations of diffusion from these molecular weights and the axon morphology support this assumption (Supplemental Fig 1). To illustrate this, Alexa 594 hydrazide and Fluo-4 fluorescence (Fig 2Aa) were normalized to their values when the first varicosity image (inset 1) was obtained. For 70 mins (until Fig 2Aa inset 2), fluorescence ratios between the dyes remained constant. Resting [Ca^2+^]_i_ was calculated from these data and the Fmax of Fluo-4 fluorescence (obtained by repetitive stimulation), applied to equation 3 (methods). [Ca^2+^]_i_ remained stable for >1 hour as dye concentrations rose (Fig 2Ab; mean resting [Ca^2+^]_i_ = 81± 5 nM; 11 cells). At 80 mins, Fluo-4 fluorescence increased more rapidly than Alexa 594’s revealing an increase in resting [Ca^2+^]_i_ (Fig 2Aa, green circles). No data were used after this.

**Figure 1.**
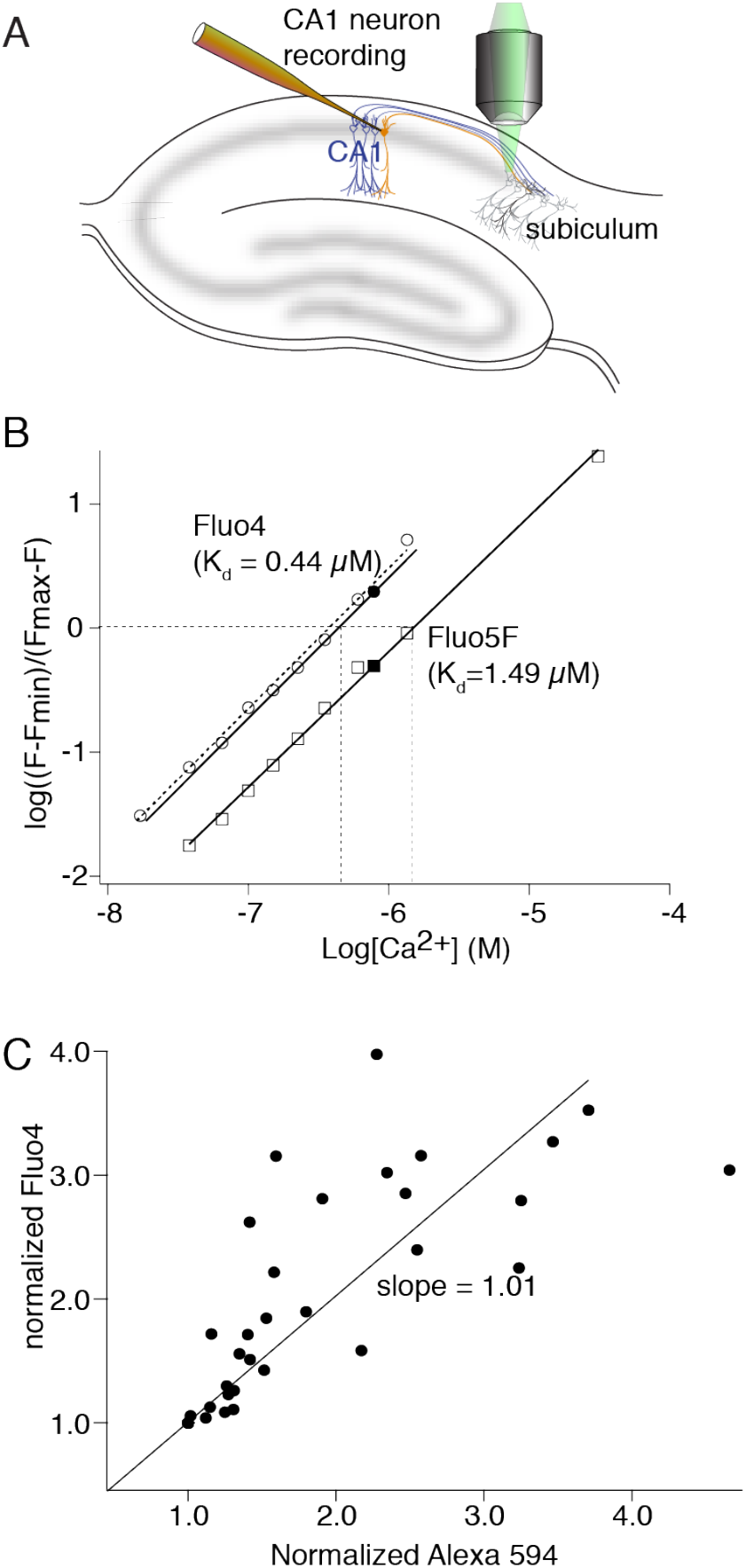
Calibration of dye diffusion and Ca^2+^ response. **A**) Recording arrangement for imaging presynaptic varicosities. CA1 pyramidal neurons were whole-cell patch clamped with electrodes containing Fluo-4 (or 5F) and Alexa 594 hydrazide. After allowing time for diffusion axons and varicosities were traced to the subiculum by imaging Alexa 594 hydrazide and Ca^2+^ was imaged using the Fluo dye. **B**) Calibration of the Ca^2+^-sensitive dyes. Fluo-4 (circles) and Fluo-5F (squares) were imaged on the confocal system used for all measurements. Fluorescence intensity was measured over a range of Ca^2+^ standard concentrations in fixed concentrations of EGTA. Log-log plots gave a slope (Hill coefficient) of 1. Ca^2+^ concentrations were applied to a patch-clamped cell by application of ionomycin. From saturated Ca^2+^ concentrations, zero Ca^2+^ and a fixed value of Ca^2+^ (0.78 *μ*M in EGTA buffered Ca^2+^ solution), the closed circle (Fluo-4) and closed square (Fluo-5F) values in the graphs were calculated. These values were used to correct the plots to values in the intracellular environment (solid lines) and to calculate values of Kd for the two dyes in the cells (0.44 and 1.49 *μ*M). **C**) The dyes introduced from the patch pipette showed equal rates of diffusion into axon varicosities. Intensities of Fluo-4 and Alexa 594 hydrazide measured by separate illumination at 488 nm and 568 nm respectively were normalized to the value of the fluorescence value obtained following identification of a varicosity and plotted against one another for points measured over the next 20 mins. The slope of a line fitted to this data through (0,0, because both dye concentrations were zero at the experiment start) is close to unity.

**Figure 2.**
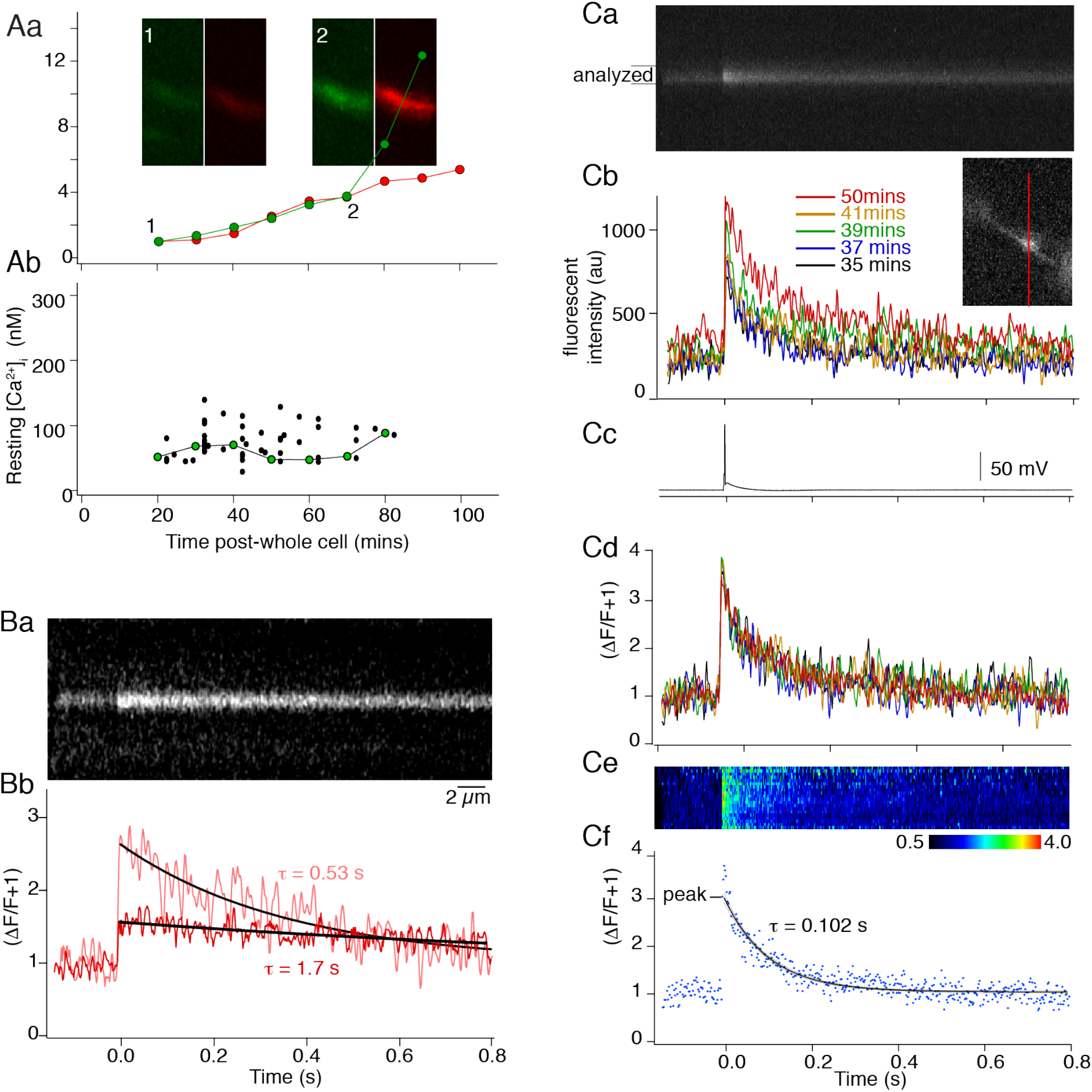
Effects of high- and low-affinity dyes on presynaptic Ca^2+^ transients. **Aa**) CA1 pyramidal neurons were recorded in whole-cell and filled with Alexa 594 hydrazide (red, 250 *μ*M) and Fluo-4 (green, 1 mM) from the pipette. Each dye’s intensity normalized to values at 20 mins post-whole cell was plotted against time in one varicosity. Insets – green, Fluo-4 and red, Alexa 594 in the same varicosity after 20 mins and 70 mins. (**Ab**) Resting [Ca^2+^] for all cells calculated from equations (1) and (2). Linked green circles – values obtained from cell in (**Aa**). **Ba**) Line scans were performed through varicosities of neurons filled with Alexa 594 hydrazide and Fluo-4, 1mM. Ca^2+^ transients were evoked by an action potential evoked at the soma. Fluo-4 was imaged during the line scan. A first action potential-evoked Ca^2+^ transient recording (∼20 mins post-whole cell) was made as dye diffused into the varicosity (transient decay rate, τ= 0.53 s, **Bb** pink). 25 mins later the peak amplitude was reduced and τ increased (τ = 1.7 s, red). **Ca**) Line scans from action-potential-evoked Ca^2+^ transient from a presynaptic varicosity (inset to **Cb**) filled with the lower affinity Ca^2+^ dye, Fluo-5F. Line scan was through the red bar. (**Cb**) Intensities of five line scans from the analyzed region in (Ca). Colors represent indicated time post whole-cell access when responses were measured. (**Cc**) Depolarizing somatic current-evoked action potentials, which evoked the Ca^2+^ transients. **Cd**) Responses from (Cb) expressed as △F/F+1 where F is the pre-stimulus Fluo-5F fluorescence. The Ca^2+^ transient is identical for each stimulus. (**Ce**) Color-coded representation of the line-scan as △F/F+1 after background subtraction from analyzed region in (Ca). (**Cf**) Single Ca^2+^ transient signal after 37 mins whole cell. The data was fit by a single exponential.

### Dependency of the Ca^2+^ signal on dye buffering capacity (κ_dye_)

Single CA1 axon varicosities loaded with high-affinity Ca^2+^-sensitive dye (Fluo-4, Kd, 0.44 *μ*M; 1 mM) were imaged by line-scanning single action potential evoked Ca^2+^ transients (Fig 2B). Transient peak amplitudes reduced, and decay time constants (τ) increased as dye concentrations rose (Fig 2B). This is consistent with Ca^2+^ buffering by Fluo-4, because the amplitude represents a proportion of the dye that is Ca^2+^-bound. As dye concentrations rise a smaller dye fraction binds Ca^2+^ entering. Unbound dye competes for Ca^2+^ with endogenous buffers, consequently, τ increases as rebinding to dye becomes more likely in cells (Neher and Augustine, 1992) or nerve terminals (Koester and Sakmann, 2000; Jackson and Redman, 2003; Brenowitz and Regehr, 2007). An alternative explanation for increased τ is diffusional loss of endogenous buffers during whole cell recording (Müller et al., 2007). However, this would occur regardless of dye concentration or affinity, be dependent on recording duration, and accompanied by increased peak amplitudes. These did not occur (supplemental Fig 2).

Similar experiments were performed substituting a low affinity Ca^2+^ dye (200 *μ*M Fluo-5F, k_d_ 1.49 *μ*M; n = 30). Responses were recorded during rising dye concentrations (Fig 2Ca-b). Normalized as (ΔF/F+1), these Ca^2+^ transients were invariant in amplitude and τ throughout the experiment (Fig. 2Cd), and the fluorescence transient was uniform across the varicosity within 6 ms of the stimulus (Fig 2Ce). Means of transients from sequential stimuli were plotted, and single exponentials fit to decays (Fig 2Cf). In all cells, transients from different varicosities at different axon locations gave reproducible amplitudes and τ’s (mean Δ[Ca^2+^]_i_ = 677 ± 10 nM (methods, equation 3), mean τ = 119 ± 1.4 ms, n = 30).

Comparing Ca^2+^ transient τ’s and amplitudes from Fluo-4 (n=11) vs. Fluo-5F (n = 7) at similar times after whole-cell access also indicates dye buffering (increasing κ_dye_) not loss of endogenous buffer causes the changes in peak ΔF/F and τ. Using Fluo-4, as concentration increased, τ increased and peak amplitudes were reduced. However, Fluo-5F left both unaltered (Supplementary Fig 2). Calculated dye concentrations were therefore used to relate dye buffering to varicosity Ca^2+^ transients (example cell, Fig 3: Supplementary materials).

**Figure 3.**
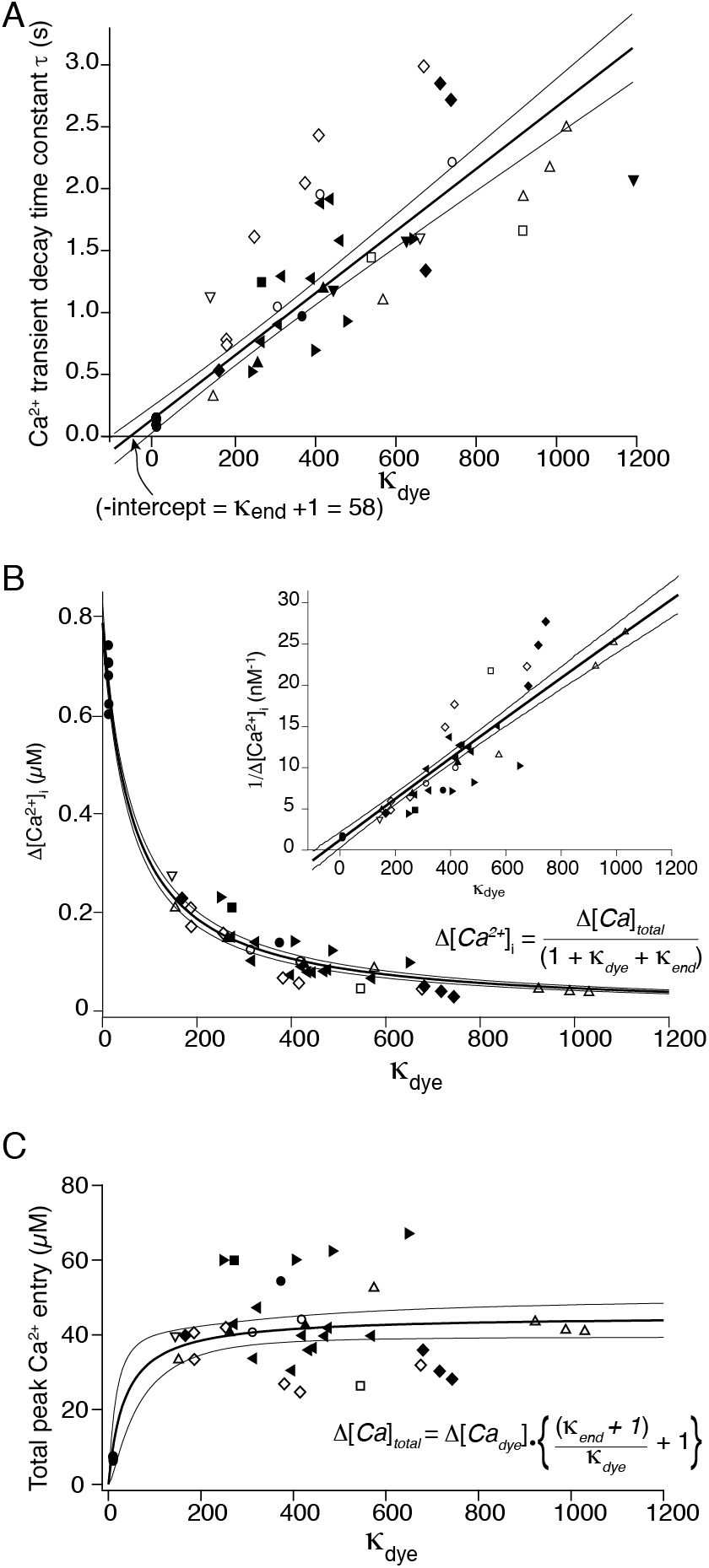
Analysis of Ca^2+^ transients and buffering using Ca^2+^-sensitive dyes. **A**) From the fits to Ca^2+^ transients obtained with Fluo-4 and Fluo-5F at various values of κ_dye_, decay time constants (τ) were plotted against κ_dye_. (Each symbol used represent a different cell) The negative intercept on the abscissa is a measure of endogenous buffering capacity (κ_end_) of the terminal (from equation 5). (**B**) Evoked change in [Ca^2+^]_i_, calculated from equation 3 is plotted against dye buffering capacity (κ_dye_) for the same data. The curved line represents a fit of equation 9. The inset represents the equivalent linear fit to the inverse of Δ[Ca^2+^]_i_, where the convergence of the fit with the abscissa represents κ_end_ and the slope is a function of Δ[Ca^2+^]_total_. The fits give values for the peak free Ca^2+^ concentration, the varicosity κ_end_ and total Ca^2+^ entering. **C**) Values of total peak [Ca^2+^] bound to dye calculated from the product of measured Δ[Ca^2+^] and the κ_dye_ were plotted against kye. These data were fit to equation 10. As the curve reaches an asymptote at high values of κ_dye_, this allows calculation of the total molar quantity of Ca^2+^ entering the varicosity following one action potential. Error bands are for 90% confidence intervals.

We determined κ_dye_ (equation 4) for each Ca^2+^ transient from varicosity dye concentrations, changes in varicosity free Ca^2+^ (Δ[Ca^2+^]i) calculated from this data, and equation (3). In 18 neurons (7 - Fluo-5F, 11 - Fluo-4) a linear fit to τ vs κ_dye_ (Fig 3A) gave an x intercept of −58 ± 40 and an estimate of the endogenous buffering capacity of the varicosity (kend) of 57 (equation 5). A plot of Δ[Ca^2+^]_i_ (equation 3) against κ_dye_ (Fig 3B) demonstrated the relationship between κ_dye_ and the Ca^2+^ transient. A fit of equation (9) to this data or a linear fit to 1/Δ[Ca^2+^]_i_ vs κ_dye_ (inset Fig 3B) gives a peak free Ca^2+^ concentration ([Ca^2+^]ι) in the absence of dye throughout the terminal of 0.76 ± 0.03 *μ*M. This is obtained from the y-axis intercept of the fit where κ_dye_ = 0. Constants from the same fit give total Ca^2+^ entry of 58 ± 6 *μ*M and κ_end_ of 75 ± 6.

Total dye bound Ca^2+^ ([*Ca_dye_*]) was calculated for each transient either from the product of [Ca^2+^]_i_ and κ_dye_, or from proportions of dye bound to Ca^2+^ calculated from the Hill equation. These gave values that differed by less than 5%. Data in Fig 3C are from the former. If total Ca^2+^ entering each varicosity were constant for each action potential, then these data are represented by equation (10), in which the asymptotic value of total Ca^2+^ entering the varicosity was 45 ± 3 *μ*M. From this and each measured varicosity dimensions (volume mean = 1.7 ± 0.3 *μ*m^3^, median = 1.2 *μ*m^3^, n=18) we determined total molar Ca^2+^ entering the varicosity (Table 1). Ke⊓d is taken from fits of equation (9) to Δ[Ca^2+^]_i_ (Fig 3B). Peak [Ca^2+^]_i_ is for Ca^2+^ throughout the varicosity and total [Ca^2+^] entering is from fits to equation (10) of data in figure 3C. Results with least errors were used and summarized in table 1.

**Table 1.** parameters of the varicosity and Ca^2+^ signaling

|  |  |
| --- | --- |
| $\kappa_{\text{end}}$ | $75 \pm 10$ |
| Resting $[Ca^{2+}]$ (nM) | $81 \pm 5$ |
| Peak $\Delta[Ca^{2+}]_i$ (nM) | $760 \pm 30$ |
| Total calcium entering ($\Delta[Ca]_{total}$) ($\mu M$) | $45 \pm 3$ |
| Total Molar $Ca^{2+}$ entry (mol) | $8.0 \times 10^{-20}$ |
| Median varicosity volume ($\mu m^3$) | $1.2 \pm 0.3$ |
| Mean varicosity surface area ($\mu m^2$) | $7.6 \pm 1.0$ |
| Fluo-5F transient decay rate $\tau$ (ms) | $119.5 \pm 1.4$ |
| Calculated $\tau$ with 0 [dye] | $0.11 \pm 0.09$ |
| Extrusion pump rate | $7.6 \times 10^6 \pm 1.1 \times 10^6 \times (\text{total } \#Ca^{2+} \text{ ions}) M^{-1} \cdot s^{-1}$ |

### Ca^2+^ source and removal from the terminal

Internal stores might contribute to Ca^2+^ transients (Cochilla and Alford, 1998; Emptage et al., 2001; Scott and Rusakov, 2008). Calculations of Ca^2+^ entering, and κ_end_ will be distorted if secondary Ca^2+^ sources exist. In recordings with Fluo-5F, ryanodine (5 *μ*M; to block store release) was superfused, and five action potentials (50 Hz; Fig. 4A) evoked a response on which ryanodine had no effect (to 111 ± 12% of control, 95% confidence interval of 90 to 132 %, n=5). Thus ryanodine does not meaningfully alter Ca^2+^ transients.

**Figure 4.**
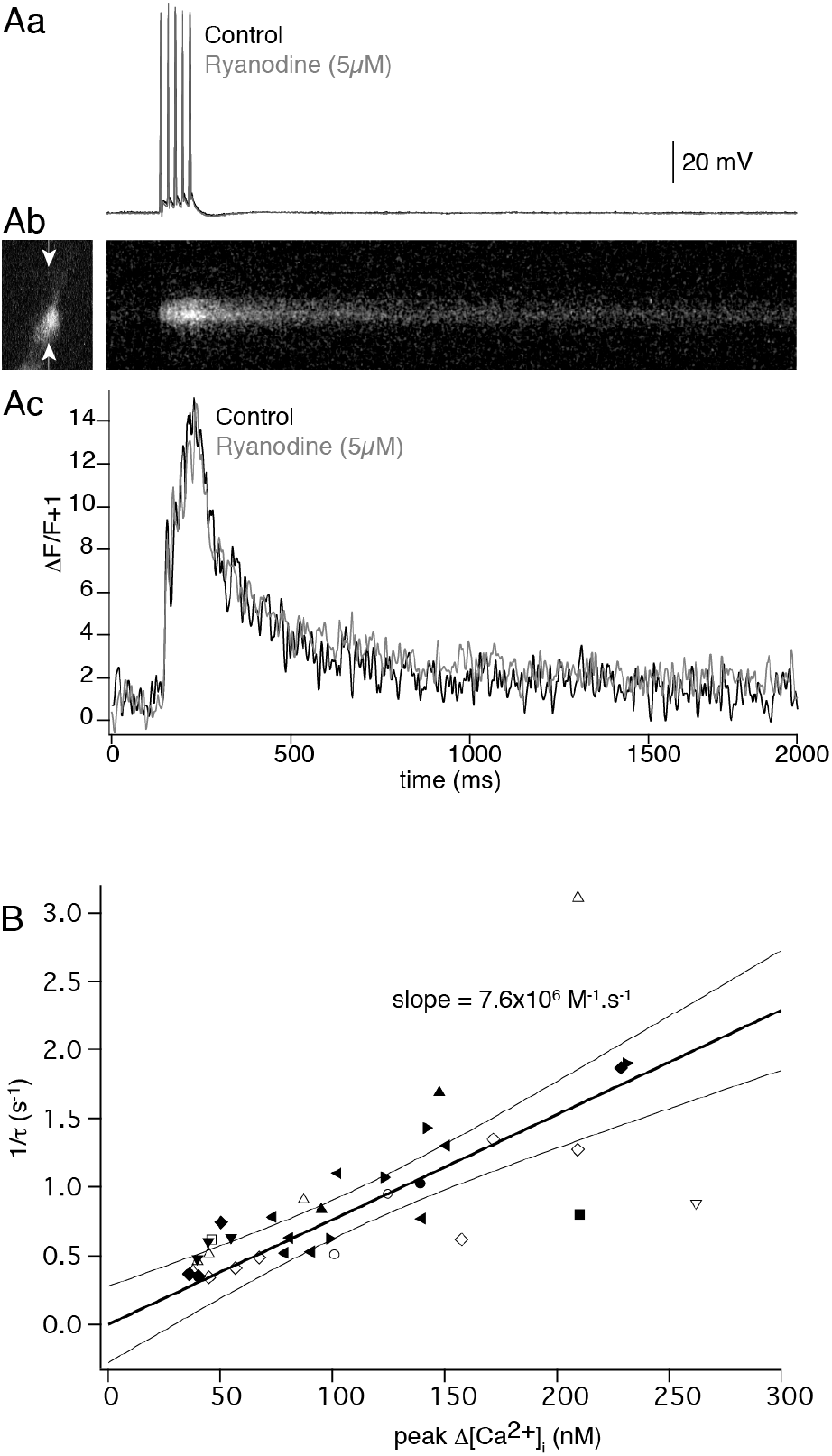
The Ca^2+^ transient is unaffected by block of release of Ca^2+^ from internal stores, and its peak amplitude is correlated with the inverse of its decay. **A**) Repetitive stimulation evokes summating Ca^2+^ transients that are unaffected by ryanodine. (**Aa**) Five action potentials at 20 Hz evokes a summating Ca^2+^ transient measured using Fluo-5F in a CA1 neuron varicosity (**Ab**) which was unaffected by a 30 min application of ryanodine (5 *μ*M; **Ac**, grey.) **B**) Inverse of the Ca^2+^ transient decay rate vs. peak Δ[Ca^2+^]; for varicosities in which the value of kjye exceeded 150. From the slope, the extrusion rate of Ca^2+^ from the varicosity was determined (equal to slope x total molar Ca^2+^ entry). (Each symbol used represent a different cell)

If Ca^2+^ extrusion is mediated by pumps with linear rates vs peak [Ca^2+^]I, then as the transient varies with κ_dye_, removal rates can be calculated. As dye concentration increases, κ_dye_ dominates κ_end_, and peak [Ca^2+^]_i_ available to pumps is reduced. Removal will be inversely proportional to available [Ca^2+^]i. The slope of 1/τ of the transient against peak [Ca^2+^]_i_ allows calculation of Ca^2+^ removal. Thus, these data obtained with Fluo-4 were plotted (Fig. 4B) and the extrusion rate (7.6 × 10^6^ × (total #Ca^2+^ ions) M^−1^·s^−1^) calculated from the slope. The linearity of these data also provide evidence that secondary Ca^2+^ sources do not contribute to the transient, at least at these values of κ_dye_. A summary of experimentally determined properties of the varicosity is given (Table 1).

### Modelling of Ca^2+^ transients in varicosities

Imaging experiments provided resting [Ca^2+^]_i_, varicosity volumes, their endogenous buffering capacity, the peak free [Ca^2+^]_i_, the total Ca^2+^ entering, and its removal rate. The principal Ca^2+^ buffer in CA1 pyramidal somata is calbindin_28K_ (Müller et al., 2005)(40 *μ*M), making it a candidate buffer in varicosities (Arszovszki et al., 2014) with well-characterised Ca^2+^ binding properties (Nägerl et al., 2000). Calmodulin with similarly characterized properties (Faas et al., 2011) has also been proposed as a dominant binding protein in these neurons. We constructed a 3D model in the simulation environment MCell (see methods) (Kerr et al., 2008), to investigate Ca^2+^ entry, diffusion, buffering, and removal during action potential stimulation. Paramaters and buffers (either calbindin_28k_ or calmodulin), used in this model were determined in this study, obtained from the literature, or varied to obtain best fits to the data (Table 2).

**Table 2.**
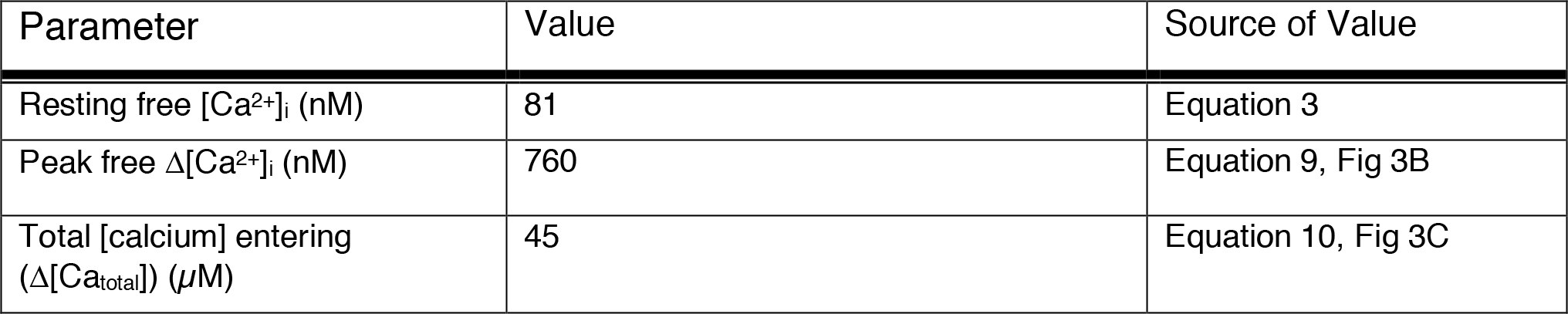
parameters used in the MCell model

A 3D mesh model was developed (Blender), from published data and from this study. Varicosities were represented by ellipsoids, 2 by 1 *μ*m (volume = 1.2×10^−11^ L, the median measured varicosity volume) *en-passant* to an axon (0.12 *μ*m diameter). They contained 320 40 nm diameter vesicles and an internal structure mimicking organelles to position 1/3 of the Ca^2+^ extrusion pumps (Fig 5A; supplementary video 1 for Ca^2+^ transient simulation). We modeled Ca^2+^ extrusion with a rate of 2.2×10^11^ M^−1^·s^−1^ distributed to 3000 pumps over the plasma membrane and organelle meshes. A Ca^2+^ leak (1.77×10^5^ s^−1^) similarly distributed achieved resting free [Ca^2+^]_i_ of 81 nM. Ca^2+^ buffering was modeled with three models of buffers (calbindin_28K_ 3:1 ratio, calbindin_28K_ 2:2 ratio or calmodulin). Calbindin28K possesses four Ca^2+^ binding sites and models have been proposed for its Ca^2+^ binding (Nägerl et al., 2000) with 3:1 or 2:2 ratios of high and medium affinity non-cooperative sites. The former model was slightly but not significantly favored in data fits *in vitro* (Nägerl et al., 2000), but the latter significantly in a model of cerebellar Purkinje neurons (Schmidt et al., 2012) in which a correction to the on-rate accounted for intracellular Mg^2+^. Ca^2+^-calmodulin binding was also modeled using published parameters (Faas et al., 2011) (table 2). We utilized the same Mg^2+^ on-rate correction in our models for calbindin_28K_ and calmodulin. Evoked Ca^2+^ entry was modeled as a total of 45 *μ*M Ca^2+^ entering the terminal in 2 ms (table 1)

**Figure 5.**
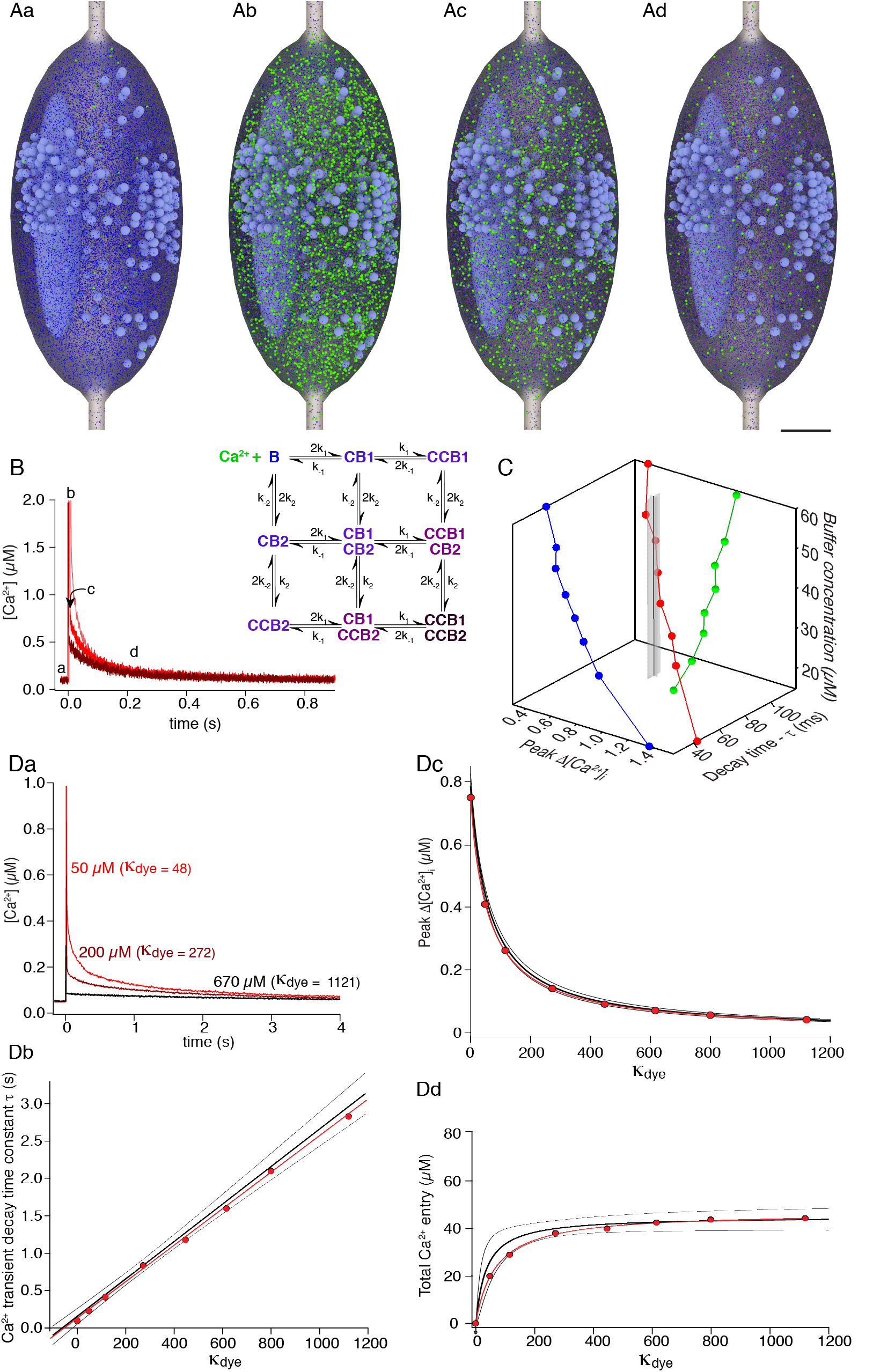
Monte Carlo simulation of Ca^2+^ transients and validation against experimental data. **A**) The varicosity was modeled as an ellipsoid 2 × 1 x1 *μ*m and contained 320 synaptic vesicles, and one large internal structure to provide sites for Ca^2+^ intrusion (i.e., ER/mitchondrion). A variable quantity of Ca^2+^ buffer (red in Aa) was modeled. Images show free Ca^2+^ ions (green) and Ca^2+^-bound states of calbindin_28K_ as per the color scheme in the kinetic model below. Times indicated by letters in B. Parameter values are from table 1 or from varying buffer concentrations. (**Aa**) pre-stimulus state (**Ab**) at peak of stimulus (**Ac**) at first time point resolveable with fluorescence imaging (Ad) after 1 s post stimulus. **B**) Simulated total varicosity free [Ca^2+^]_i_, following stimuli at three calbindin_28k_ concentrations (15 *μ*M, pink; 40 *μ*M, red; 60 *μ*M, black). Single exponential fits (red) were applied to these simulations and values of peak Δ[Ca^2+^] and decay (τ) were obtained from these fits. Inset shows kinetic scheme for the calbindin_28k_ 2:2 ratio. Parameters in table 2. **C**) 3D plot of simulated τ, peak ΔCa^2+^ against varying buffer concentrations (15 – 60 *μ*M) following simulated varicosity stimulation. Blue – 3:1 ratio model of calbindin_28k_, red – 2:2 model of calbindin_28K_, green – model of calmodulin. Model parameters in table 2. The vertical black line and gray shading represent the values of experimentally obtained τ, peak ΔCa^2+^ and associated standard errors. These data converge only with the 2:2 ratio of calbindin_28k_ at a concentration of 39.7 *μ*M. **D**) Varying Ca^2+^-sensitive dye concentrations (Fluo-4, see table 1) were simulated and resultant Ca^2+^ transients graphed. (**Da**) Responses shown with dye concentrations of 50, 200, and 670 *μ*M. (**Db-Dd**) decay time, peak ΔCa^2+^, and total Ca^2+^ entry from these simulated transients over values of κ_dye_ from 0 to 1121 (red circles and lines) is plotted as per Fig 3C and overlaid on the fits and 90% confidence intervals of that experimental data.

Buffer concentrations were first estimated by solving the Hill equation for values of total Ca^2+^ entry, and resting, and stimulated peak free [Ca^2+^]_i_. This assumes equilibrium at peak, and was expected to underestimate true buffer concentrations because Ca^2+^ transients are too rapid for equilibration. This gave 92.8 *μ*M for calmodulin and 24.8 *μ*M calbindin_28K_ (2:2 ratio of high and medium affinity sites), or 19.7 *μ*M (3:1 ratio).

To determine buffer parameters, simulated endogenous buffer concentrations were varied starting at steady-state results to compare simulations to experimentally determined peak free Δ[Ca^2+^]_i_ and τ - (Fig 5B; parameters - Tables 2,3). Transient decays were fit with single exponentials omitting the first 4 ms of the simulation to avoid initial Ca^2+^ inhomogeneities. From these fits, peak free Δ[Ca^2+^]_i_ and τ were plotted against simulated buffer concentrations (Fig. 5C). Intersection of simulated and experimental values for τ and Δ[Ca^2+^]_i_ implies that model parameters are accurate and provides an estimate for buffer concentration and type. Models of calbindin_28K_ with a 2:2 binding ratio converge to within the 95% confidence interval of the experimental data (Fig. 5C). A least squares fit for this convergence predicts a calbindin_28K_ 2:2 concentration of 39.7 *μ*M – in close agreement to 40-45 *μ*M calbindin_28K_ obtained experimentally for rat CA1 pyramidal neuron somata (Müller et al., 2005). Neither the calbindin_28K_ 3:1 ratio nor calmodulin converged (Fig 5C, supplementary Fig 4). Note, in cerebellar purkinje neurons, models for calbindin_28K_ also favor 2:2 ratios (Schmidt et al., 2012). While other Ca^2+^ buffers are present, we conclude that simulating calbindin_28K_ with a 2:2 ratio of binding site has validity.

**Table 3.** MCell model specific parameters.

|  |  |
| --- | --- |
| Iteration interval (s) | 4x10<sup>-8</sup> (calbindin/calmodulin buffer determination).<br>5x10<sup>-9</sup> (syt1 and local $\text{Ca}^{2+}$) |
| Partition size ($\mu\text{m}$) (this is an MCell function that divides the volume into subvolumes to optimize calculations) | 0.2 |
| Molecular interaction radius (nm) (radius at which two molecules may interact) | 10 |
| Microscopic reversibility (function that increase accuracy of reactions) | on |

### Model validation of experimental results in a small terminal

We used simulations to determine if our experimental approach is valid for small terminals. Effects of Fluo-4 (0 to 670 *μ*M) were simulated and κ_dye_ calculated as for experimental data. As for experimental data, rising values of simulated κ_dye_ reduced peak [Ca^2+^]_i_ and increased τ. Peak [Ca^2+^]_i_ and τ were measured from single exponentials fitted to simulated data (Fig. 5Da) ommitting the initial 4 ms. Results were plotted with experimental data as τ vs. κ_dye_ (Fig. 5Db), peak [Ca^2+^]_i_ vs. κ_dye_ (Fig. 5Dc), and total Ca^2+^ entry captured by the dye vs κ_dye_ (Fig. 5Dd). In all cases simulations fell within the 90% confidence limits of experimental data supporting the use of this approach in small varicosities.

We then compared experimental Fluo-5F results to simulations of Fluo-5F-Ca^2+^ binding to validate simulations with experimental data. The simulation reproduced experimental measurements showing that Ca^2+^ transients remain isolated to varicosities (supplemental Fig 5A,B). Additionally, simulated responses to repetitive stimulation fell within the 90% confidence interval of the experimental data to equivalent stimuli (supplementary Fig 5C).

### Spatio-temporal distribution of Ca^2+^ entry to the terminal

Calculations of varicosity [Ca^2+^] and its buffering assume that Ca^2+^ rapidly reaches spatial uniformity within varicosities. The overlap between single exponential fits to experimental data and fits to simulations indicate this approach is valid to calculate total Ca^2+^ entry and its buffering. However, in five neurons recorded with sufficiently high-resolution, we observed repeatable, but non-uniform Ca^2+^ distributions immediately after stimulation (Fig 6).

**Figure 6.**
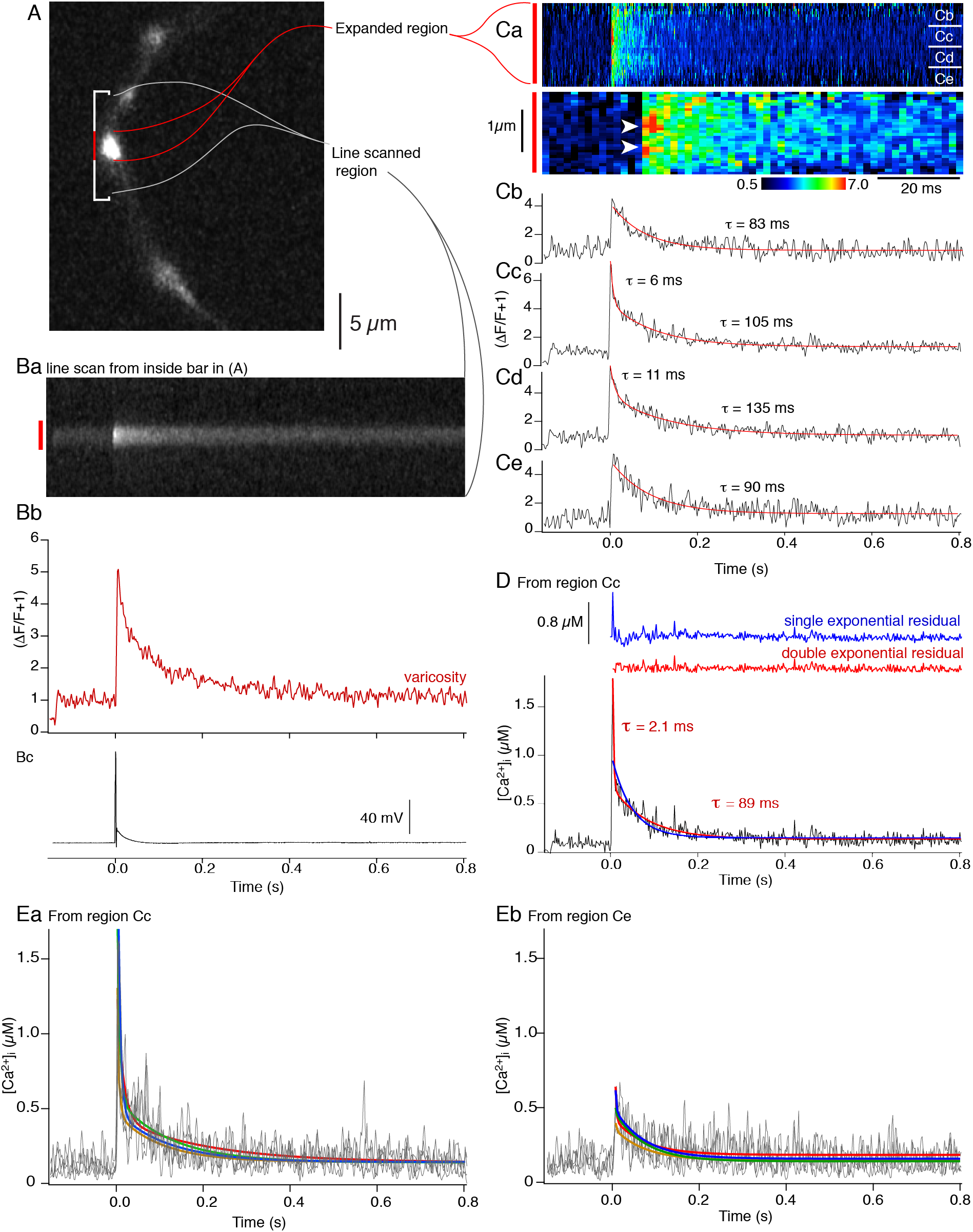
Ca^2+^ or Ca^2+^ dye complex can show rapid diffusion from presynaptic terminal hotspots immediately after the stimulus. **A**) Axon varicosity containing both Alexa 594 hydrazide and Fluo-5F imaged using the Alexa dye. Bracketed region encompasses a single varicosity (center, red) and axon on either side (white) used for line scanning. **B**) Mean of 4 line scans from region in (**A**). The varicosity exhibited a Ca^2+^ transient similar to those recorded in earlier figures (**Ba**). Mean intensity of the linescan through the varicosity (**Bb, red bar**) which showed a large increase in Ca^2+^ dye fluorescence following a single action potential (**Bc**), **C**) The mean of 4 line-scans within the varicosity (red in **B**) displayed as ΔF/F+1 (**Ca**, upper panel time scale as in graphs in D and E below). Immediately after the stimulus there is a rapidly decaying component of the transient. The lower panel of **Ca** shows the two hotspots (arrowheads) at a 10x timescale. Data was then analyzed in quadrants (**Cb-Ce**). Center quadrants (**Cc and Cd**). The center quadrants were well fit with double exponentials, the outer with single exponentials. **D**) Data in **Cc** replotted to demonstrate the amplitude of free [Ca^2+^] within this region of the varicosity. As with the data in Cd, this required a double exponential fit (single exponential and residual in blue, double exponential and residual in red). **E**) Ca^2+^ transients (grey) were plotted for each of the 4 sequential responses averaged in (C). (**Ea**) plotted from the region labeled (Cc) and (**Eb**) from the region labeled (Ce) in (Ca). Double exponentials were fit to these data for each of the four traces (colored traces).

In each varicosity (Fig. 6A), line-scans (mean of 4 transients in each of the 5 terminals; ΔF/F+1 vs time; Fig. 6B), reveal brighter regions immediately post-stimulus (Fig. 6Ca, hotspots arrowed at faster timebase - lower panel). However, these spots are close to the resolution limit, and their intensity might be affected by errors in background or prestimulus intensity. If hotspots represent localized Ca^2+^ entry then a faster local decay would represent diffusion from this site (mean τ overall = 119.5 ± 1.4 ms). Therefore, τ was analyzed in line-scan subregions. In the case illustrated, a single exponential well-fit Ca^2+^ transients away from hotspots (Fig 6Cb,e; sum of squares of residuals were not significantly different between single and double exponential fits, p = 0.11) but did not adequately fit regions at hotspots. These were well-fit by double exponentials (τ’s of 6 - 11 and 105 - 135 ms, Fig 6Cc,d). Similar results were obtained in all 5 neurons (mean τ1 = 9.0 ± 2.9 ms; τ2 = 124.5 ± 19.1 ms;; sum of squares of residuals were significantly different between single and double exponential fits at these hotspots, p = 0.013). To illustrate the peak [Ca^2+^]_i_ recorded by Fluo-5F, experimental data is replotted (Fig. 6D, black) as [Ca^2+^]_i_ vs. time, and is well-fit with a double exponential (red; residual above also in red, fast τ = 2.1 ms, peak [Ca^2+^]_i_ of 1.8 *μ*M; vs. 0.8 *μ*M for the rest of the varicosity). In all 5 neurons the mean peak free [Ca^2+^]_i_ = 2.7 ± 0.57 *μ*M and mean fast τ = 3.3 ± 1.3 ms. By comparison, the data was poorly fit by a single exponential (blue, residual above, goodness of fit was again determined by comparing sums of squares of residuals. These were significantly different between single and double exponential fits at hotspots; p = 0.02, but not away from hotspots, p = 0.06).

These Ca^2+^ hotspots are repeatable. Examples from two locations (arrowed Fig 6C) with an early fast Ca^2+^ transient (Fig 6Ea) or lacking one (Fig 6Eb) were analyzed in 4 sequential stimuli (5 cells). Each response was fit with a double exponential. Fast exponential amplitudes for the two regions were significantly different (p = 0.0052, two factor ANOVA), but there was no significant difference between slower exponential amplitudes (p = 0.08).

We determined whether discrete placement of Ca^2+^ entry within the simulation could reproduce the experimental Ca^2+^ distribution. In simulations, Ca^2+^ entry was located at 1 to 6 plasma membrane sites. Experimental line-scanning was simulated (20 random seeds) by simulating values of Fluo-5F ΔF/F+1 (Fig 7A) in a line of discrete volumes across the model varicosity (Fig. 7B, vertical yellow band). One end of this band always included only one Ca^2+^ entry site. ΔF/F+1 values, were resampled to rates obtained during experimental line scanning (500 Hz) to create a simulated line-scan matrix (Fig 7A) equivalent to the experimental data.

**Figure 7.**
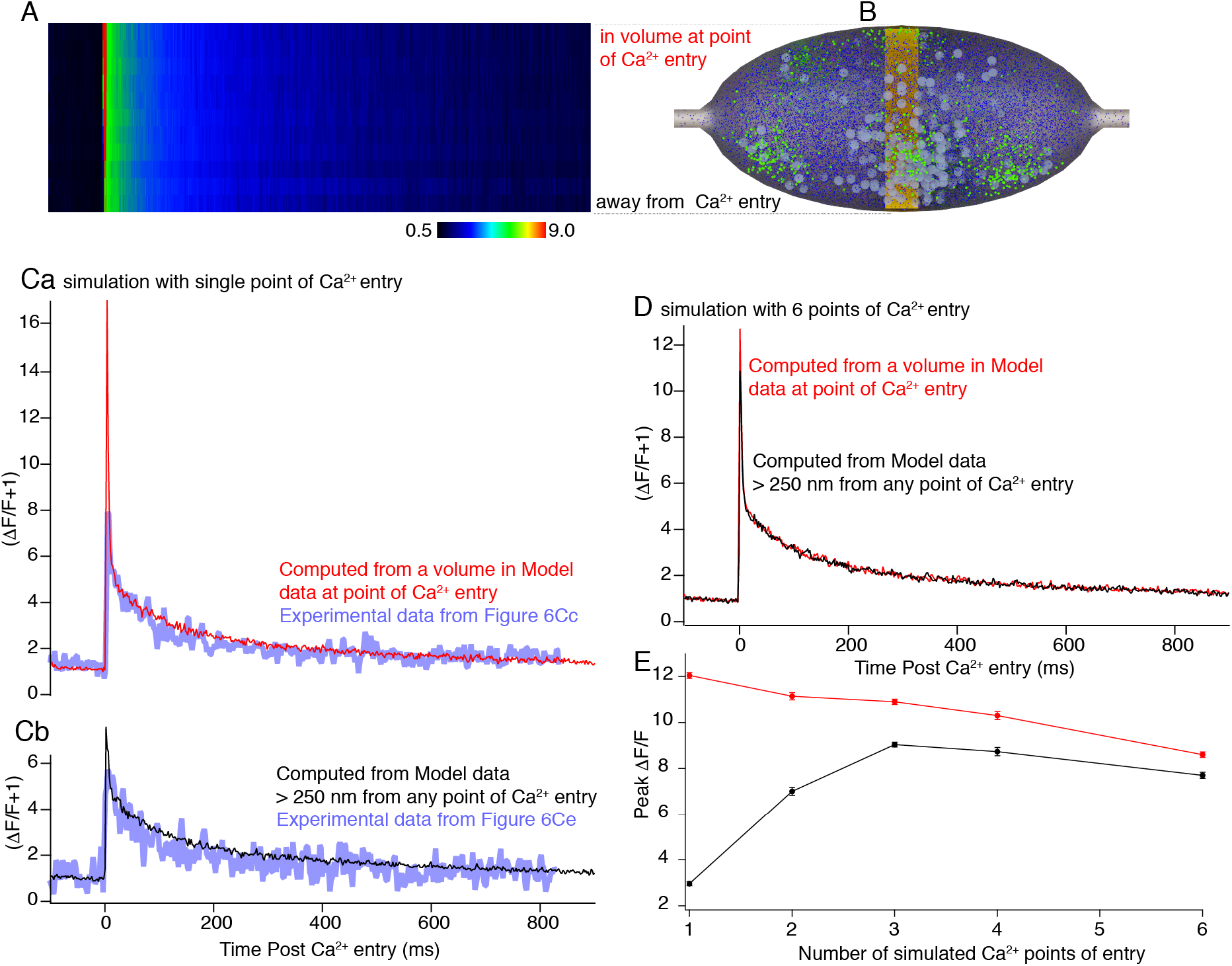
Simulations of Fluo-5F transients and clustering of Ca^2+^ entry. Line scans (**A**) of Fluo-5F transients were simulated by measuring its simulated Ca^2+^ binding in a row of volumes (yellow boxes) across the varicosity (**B**). **C**) Ca^2+^ entry was simulated at 1 discrete site placed at the plasmalemma, at the top of the yellow band in (B). (**Ca**) Simulated Fluo-5F ΔF/F+1 transient (red) in the volume closest to the site of Ca^2+^ entry (top of yellow band). Experimental data from Fig 7Cc overlaid in blue. (**Cb**) Simulated Fluo-5F ΔF/F+1 transient in the volume at the bottom of the yellow band in (B) opposite the site of Ca^2+^ entry (black). Experimental data from Fig 7Ce overlaid in blue. **D**) Ca^2+^ entry was simulated at 6 sites, including one placed at the plasmalemma, at the top of the yellow band in (B). The Fluo-5F transient is shown from this site (red) and away from sites of Ca^2+^ entry from the bottom of the yellow band in (B) (black). **E**) Graph of peak amplitude of a double exponential fit to the simulated transient at a site of Ca^2+^ entry (red) and away from these sites (black) in varicosities at which Ca^2+^ entry was at 1 to 6 sites.

Simulation results were plotted from the two ends of the yellow band, obtained when all Ca^2+^ entry (45 *μ*M) was at one point (Fig 7B, top – point of simulated Ca^2+^ entry, bottom – away from Ca^2+^ entry) (Fig 7C, overlaid with experimental data, blue). There is a substantial difference between amplitudes of the earliest peak at the Ca^2+^ entry site (7Ca, red) compared to the other side of the simulated varicosity (Fig 7Cb, black). Similar simulation results were plotted where Ca^2+^ was evenly distributed at 6 locations, one of which was at the same location as above. Little difference in peak amplitude was seen at the Ca^2+^ entry site and away from it (Fig 7D).

Double exponentials were fit to the simulations. τ’s of fast exponentials were within the 95% confidence limits of fits of early components of experimental data (simulations from 3-4 ms; experimental data, Fig 6D, 3.3 ± 1.3 ms). Peak amplitudes of simulated Fluo-5F ΔF/F early components were obtained for fits to all distributions of Ca^2+^ entry at the site of entry and across the varicosity at the opposite end of the yellow band (Fig 7B) from this site. Substantial differences in peak amplitudes at a point of Ca^2+^ entry compared to the opposite side of the varicosity away from Ca^2+^ entry, were observed only when Ca^2+^ entry was at one or two sites (that is when at least 50% of total varicosity Ca^2+^ entry was localized to one site, Fig 7E). Thus, to obtain experimental local peaks in Ca^2+^ (Fig 7), clustering of VGCCs may occur.

### Proximity of the point of Ca^2+^ entry to the release machinery and Paired-Pulse Facilitation

Proximity of Ca^2+^ entry to its molecular targets can be estimated by comparing effects of BAPTA (rapidly binds Ca^2+^) to EGTA (slower binding) (Adler et al., 1991). To record synaptic responses from CA1 synapses, their axons were stimulated (1/15 Hz) (Hamid et al., 2014). Whole-cell recordings were made from subicular pyramidal neurons and excitatory postsynaptic currents (EPSCs) recorded (in bicuculline, 5 *μ*M; AP5, 50 *μ*M) to isolate AMPA receptor responses. BAPTA-AM was superfused (10-100 *μ*M, 20 mins) and reduced EPSCs dose-dependently (to 35 ± 13 %, 100 *μ*M, n = 4; to 38 and 49%, 20 *μ*M and to 80 ± 10 % of control, 10 *μ*M, n = 3; Fig 8A). In contrast EGTA-AM (20 – 100 *μ*M was ineffective (100 *μ*M, n = 2, to 91 and 101%; 20 *μ*M, n = 7 to 110 ± 30% of control, p = 0.86, Fig 8C). These results imply a close spatial association between VGCCs and syt1 responsible for exocytosis.

**Figure 8.**
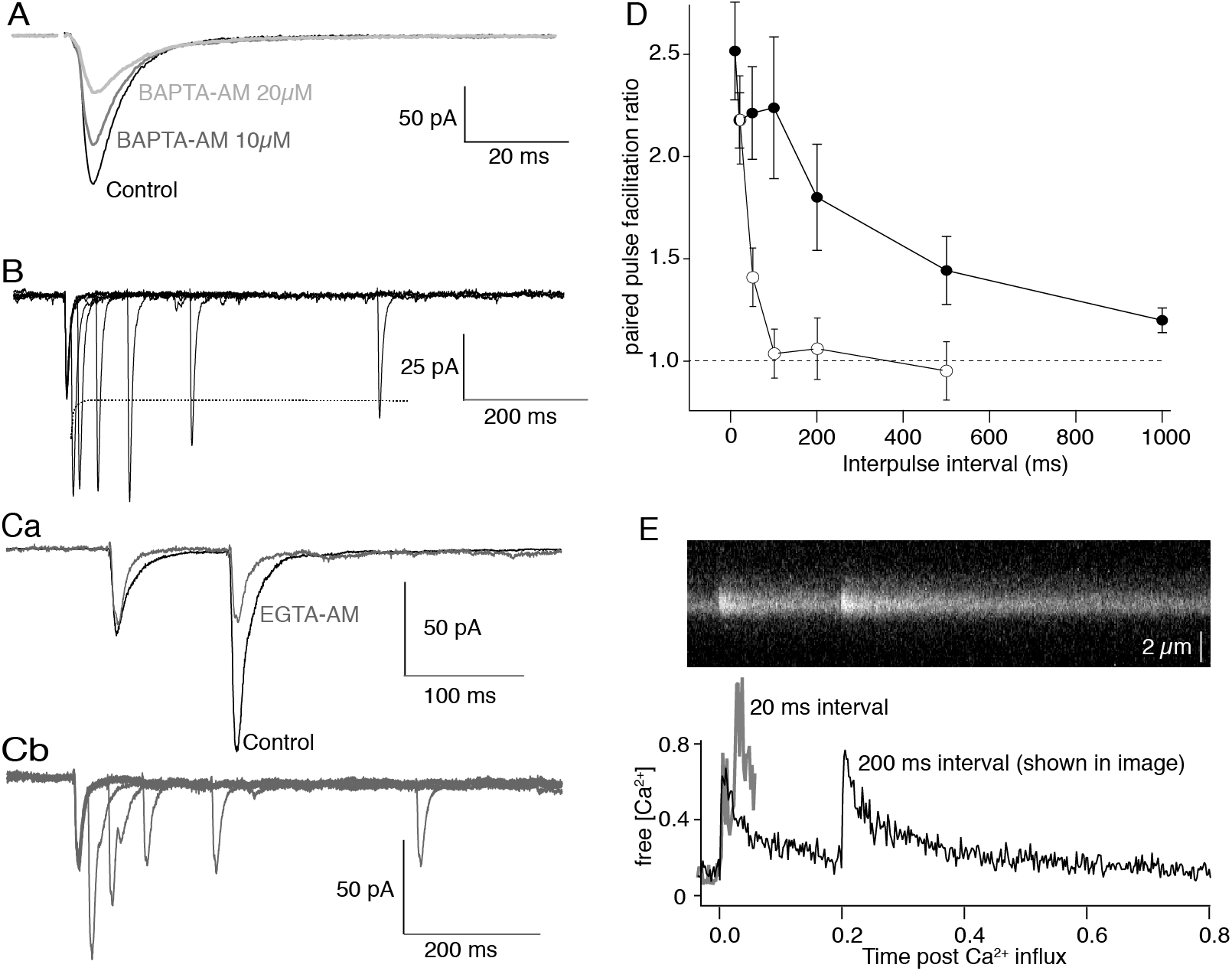
Paired-pulse potentiation is not explained by presynaptic residual Ca^2+^ concentration. **A**) EPSCs were recorded in subicular pyramidal neurons following stimulation of CA1 pyramidal axons. BAPTA-AM (10 – 20 *μ*M) was applied to the superfusate for 20 mins leading to substantial dose-dependent inhibition of the evoked EPSC. **B**) Paired-pulse plasticity in which CA1 axons were stimulated at intervals of 10 ms to 500 ms and the EPSCs recorded in subicular neurons. The dashed line represents simple summation from the first stimulus. All intervals generated potentiation. **C**) Paired-pulse responses were obtained at 100 ms interstimulus intervals from 5 neurons in which EGTA-AM (10 *μ*M) was subsequently added to the superfusate (**Ca**, grey). Paired-pulse facilitation was prevented. (**Cb**) In 5 neurons pretreated with EGTA-AM, the interstimulus interval was varied from 20 to 500 ms to demonstrate that the paired pulse potentiation was prevented over intervals greater than 50 ms. **D**) Control paired-pulse ratios against interstimulus intervals obtained from 7 neurons stimulated as in (**Ab**) (filled circles) and from 5 neurons recorded after EGTA-AM application (open circles). **E**) Ca^2+^ transients were stimulated in paired pulses. Top panel shows line-scan image of stimuli at 200 ms intervals. Bottom panel shows paired-pulse responses at 20 (grey) and 200 (black) ms intervals.

Repetitive stimulation may evoke Ca^2+^-dependent facilitation (Zucker and Regehr, 2002) caused by residual Ca^2+^, Ca^2+^-buffer saturation (Klingauf and Neher, 1997; Matveev et al., 2004), or Ca^2+^-dependent processes distinct from the exocytic machinery (Fioravante and Regehr, 2011). We determined time-courses of paired-pulse facilitation with EPSCs in subicular pyramidal neurons (Fig 8B; n=7). Intervals at 100 ms gave a mean facilitation ratio of 1.85 ± 0.191 (pulse 2/pulse 1, n=5, p = 0.014, Fig 8Ca) which was abolished by EGTA-AM (20 *μ*M; ratio after EGTA-AM = 1.0 ± 0.1, p = 0.001). In three of these cells the paired-pulse interval was varied (20 to 500 ms, Fig 8B, C,D). In EGTA, facilitation was abolished at intervals >50 ms. Facilitation ratios from controls (n = 7) and after EGTA (n = 3) were plotted from 20 to 1000 ms intervals (∼duration of varicosity Ca^2+^ transients). The effect of paired-pulse stimulation on Ca^2+^ transients was measured at 20 and 200 ms intervals. Amplitudes of the paired evoked Ca^2+^ transients (Δ[Ca^2+^]_i_) was not significantly altered (n = 5 and 3, Fig 8E; p = 0.25 at 20 ms). Thus, paired pulse facilitation is Ca^2+^-dependent, but Ca^2+^ transients do not measurably summate.

### Simulating Paired-Pulse Presynaptic Ca^2+^ transients

To address experimental limitations of analyzing Ca^2+^ at the spatio-temporal resolutions of the vesicle fusion machinery, we simulated Ca^2+^, Ca^2+^ buffer states, and effects of repetitive stimulation on free Ca^2+^ using parameters previously determined (Fig 9; in Supplementary video 2). Within the simulation, at rest, >90% of calbindin_28K_ is unbound. Stimulation causes partial occupancy of all calbindin_28k_ states (Fig 9A). However, 2/3 of all bound states remain unoccupied even at peak (Fig 9B, Supplementary Fig 6 shows all calbindin_28k_ states). Nevertheless, unbinding is slow and full recovery takes longer than 1 s. We then determined the effect of paired pulses over intervals from 20 to 1000 ms. Though calbindin_28K_ was not saturated at all intervals (Fig 9B), the second pulse achieved higher peak free [Ca^2+^] than the first (Fig 9C, difference between 2^nd^ peaks, black, and red dashed line). This is in spite of the fact that the Ca^2+^ signal recorded at the base of this initial transient resolvable with imaging is only enhanced by <200 nm and only at the very shortest intervals (summation of the component resolvable by experimental imaging is shown by the blue dashed line).

**Figure 9.**
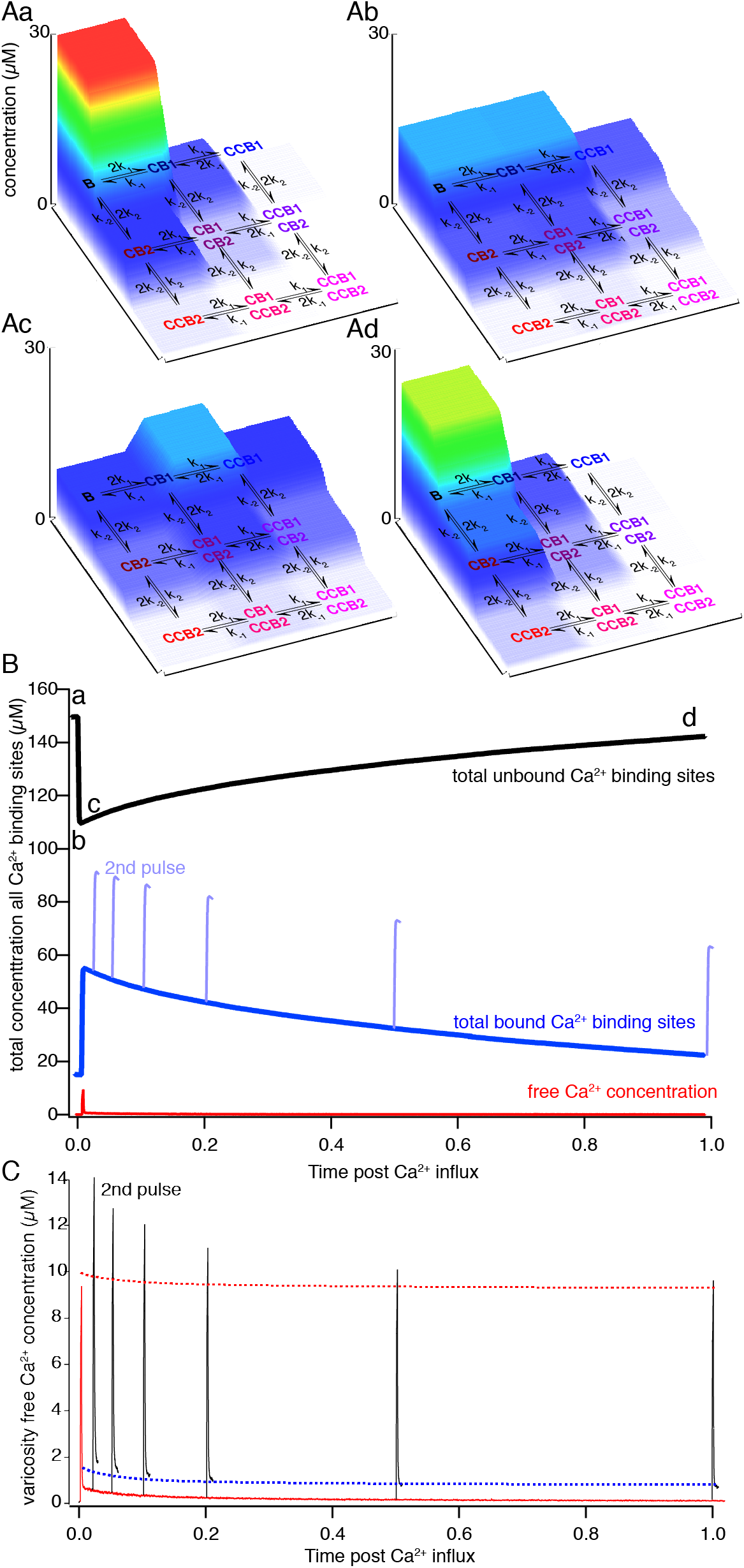
Simulations of Ca^2+^ buffering in the varicosity. **A**) Using simulation parameters previously determined, the kinetics of calbindi_28K_ binding were simulated and displayed as state diagrams from before a simulated stimulus (Aa) as well as during and after the stimulus (Ab,c and d) from time points indicated in (B). Concentrations of calbindin_28K_ in each of its possible Ca^2+^-bound or unbound states throughout the varicosity is plotted on the vertical axis and color coded to concentration. The horizontal axes display Ca^2+^ binding to high (B1)- and medium-affinity (B2) sites (C – single Ca^2+^ bound, CC – 2 Ca^2+^ bound). **B**) Graph of total concentration of vacant Ca^2+^-binding sites on calbindin_28K_ throughout the varicosity (black) and total bound Ca^2+^ (blue) after a stimulus and 2^nd^ pulse in light blue at varying interpulse intervals. Free Ca^2+^ concentration is also shown (red) to the same scale for a single pulse. **C**) Graph of free Ca^2+^ concentration after a single stimulus (black) throughout the varicosity and after 2^nd^ stimuli at varying paired-pulse intervals (black). Red dashed line indicates theoretical value of linear summation of a second pulse. Blue dashed line represents the linear sum of the Ca^2+^ signal that is resolvable using imaging at 500 Hz.

**Figure 11.**
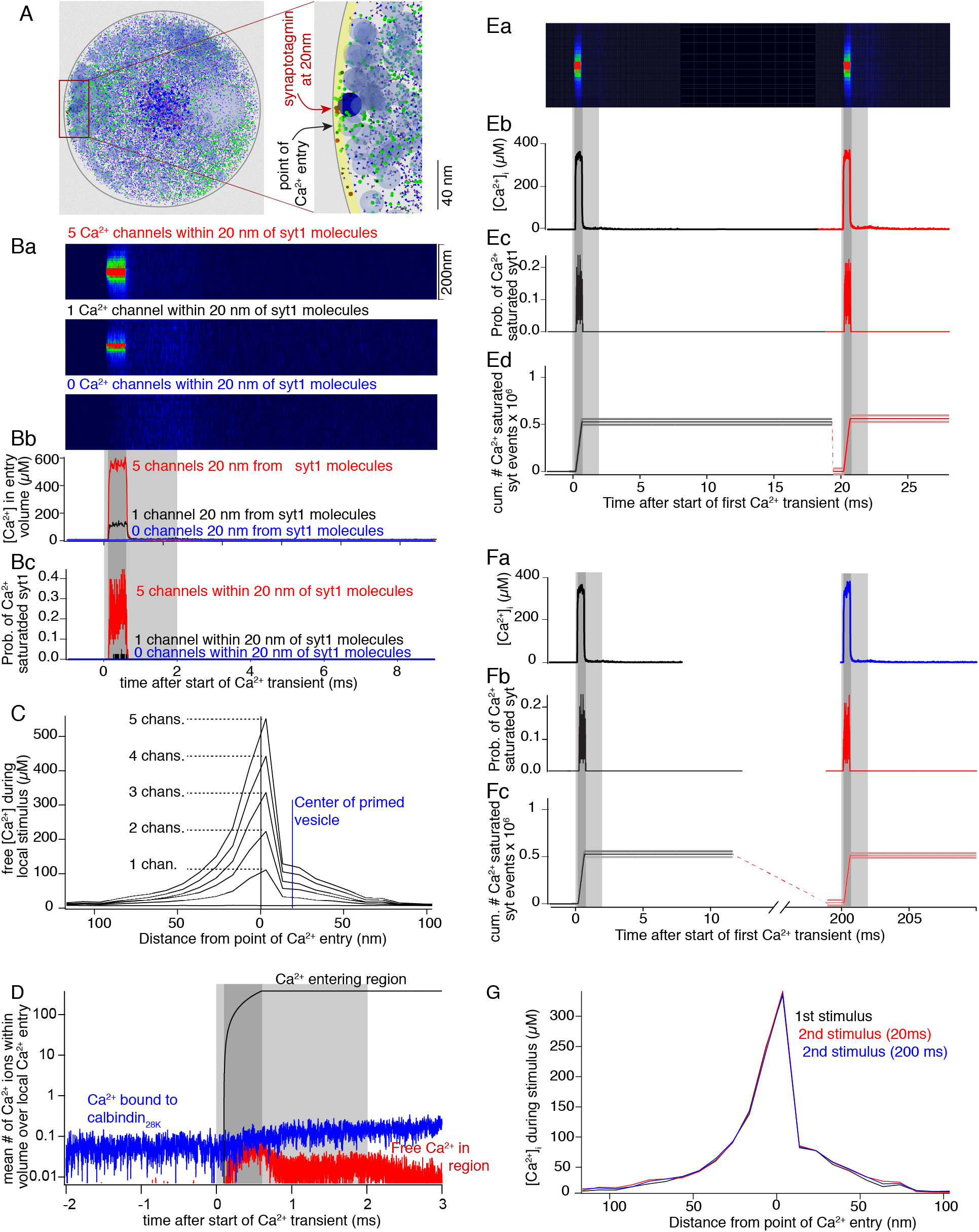
Paired pulses do not alter nanometer domains of Ca^2+^ entry. **A**) The characteristics of Ca^2+^ entry, buffering, and dispersal were analyzed close to points of Ca^2+^ entry modeled within 20 nm of a vesicle along the plasmalemma. Left hand panel shows view along the axon axis including interior contents of the varicosity (small blue dots—calbindin_28K_, green—Ca^2+^ vesicles, and light blue—intracellular organelle). The right panel is a higher magnification of the region analyzed showing an area adjacent the membrane (yellow band, where simulated Ca^2+^ was measured) and vesicles (blue circles, including one dark blue at the site of Ca^2+^ entry). It is this vesicle that distorts the spatial symmetry of the Ca^2+^ signal in D and G similar to (Shahrezaei and Delaney, 2004). **Ba**) Ca^2+^ transients expressed as a simulated line-scan over time from the yellow band in (A) centered at one point of Ca^2+^ entry for the sum of 5 channels (red), 1 channel (black) and 0 channels (blue) active for 0.5 ms (dark grey) against a background of 45 *μ*M entering the entire varicosity for 2 ms (light grey) in (**Bb**) where data is from the center 20 nm of (**Ba**). (**Bc**) Probability of synaptotagmin 1 model showing occupancy of all five Ca^2+^ binding sites for five (red) and one (black) local Ca^2+^ channels. Gray shaded regions indicate timing of Ca^2+^ entry (dark-at the local point of Ca^2+^ entry; light which also encomasses the dark grey period – from 0 to 2 ms of remaining Ca^2+^ entering elsewhere in the varicosity). **C**) Spatial distribution of the peaks of the Ca^2+^ transients from the time point at the end of the dark gray shading in (B) for 0 through 5 channels at the local site of Ca^2+^ entry. **D**) Plot of total Ca^2+^ entering a Ca^2+^ domain within 20 nm of three channels (black), the free Ca^2+^ in this region, and the Ca^2+^ in the region bound to calcium buffer. The loss of Ca^2+^ to diffusion is the difference between the black trace and the sum of the black and red traces (Note that these traces are on a logarithmic scale.) Gray shaded regions as per (B). **E**) Paired-pulse Ca^2+^ transients from the yellow region in (A) simulating three local channels located within 20 nm of synaptotagmin 1. (**Ea**) Simulated line-scans as for (**Ba**), but with a second pulse simulated 20 ms after the first. (**Eb**) Ca^2+^ transients within 20 nm of local Ca^2+^ entry repeated at a 20 ms interval. (**Ec**) Probability of synaptotagmin 1 binding five Ca^2+^ ions. (**Ed**) Cumulative number of synaptotagmin 1-five Ca^2+^-binding events to emphasize these are equal in both pulses. (**F**) Paired pulse results (**Fa-Fc**) as per (**Eb-Ed**) but at a paired-pulse interval of 200 ms. (**G**) Spatial distribution of the peak free(?) Ca^2+^ entry in pulse 1 (black) and the second pulse at 20 ms (red) and 200 ms (blue).

### Activation of synaptotagmin 1 by evoked presynaptic Ca^2+^ entry

Synaptotagmin 1 (syt1), the Ca^2+^ sensor for exocytosis, has two C2 domains with 5 Ca^2+^ binding sites, some of which have mM affinities for Ca^2+^. This requires synaptotagmin to be <100 nm of the Ca^2+^ source (Fig 8) (Adler et al., 1991; Augustine et al., 1991a) and leaves unanswered whether all sites must bind Ca^2+^ to evoke exocytosis (Radhakrishnan et al., 2009). We simulated syt1 Ca^2+^ binding at the plasmalemma (table 2) using data that membrane interaction of the C2A domain enhances its Ca^2+^ binding (Radhakrishnan et al., 2009). Ca^2+^ entry and buffering were again simulated. To determine the requirements for Ca^2+^ entry in the immediate vicinity of syt1 for full binding to occur, simulations were performed with 6 Ca^2+^ entry sources across the surface of the varicosity such that a total of 45 *μ*M entered at each stimulus. Syt1 molecules were placed at 20, 100 and 200 nm from one of these sources (Fig 10A), adjacent to a vesicle. This proximal Ca^2+^ source varied from an equivalent of 0 to 5 simulated channels (0.25 pA each, 0.5 ms open time).

More than 20 nm from the proximal Ca^2+^ source no full syt1 binding events were recorded (100 random seeded simulations). Thus, the peak free [Ca^2+^]_i_ throughout the varicosity (9.3 *μ*M; Fig 9C red) is insufficient to occupy all 5 syt1 binding sites. However, even one simulated VGCC within 20 nm of syt1 allowed this binding (Fig 10Bc, black) and more channels increased this probability (Fig 10Bc, red). With 5 channels there was a small probability of syt1 full binding 100 nm from the Ca^2+^ source. Within 20 nm from the Ca^2+^ source, one channel raised the transient concentration to 110 *μ*M; 5 channels to 560 *μ*M (Fig 10Bb).

The narrow spatial halfwidths and rapid decay of Ca^2+^ within tens of nm of VGCCs indicate rapid Ca^2+^ removal from these volumes. Plots of total Ca^2+^ entering the 20 nm scale region, Ca^2+^ bound to calbindin_28K_ within the region, and free Ca^2+^ are shown on a log scale (Fig 10D) to encompass the range of concentrations when activation of 3 VGCCs was simulated at this site (similar results were obtained by clustering all Ca^2+^ entry at these points). Calculations of the difference between Ca^2+^ entry and [Ca^2+^]_i_ and CaDye indicate that at this scale removal is dominated by diffusion. Thus, local Ca^2+^ concentrations are dominated by diffusion during paired pulse stimulation when Ca^2+^ buffers throughout the varicosity are not close to saturation (Fig 9). Consequently, no significant paired-pulse facilitation (20 and 200 ms intervals) of the local Ca^2+^ signal (within 200 nm of VGCCs) or of full syt1-Ca^2+^ binding occurred (Fig 10E,F).

## Discussion

CA1 pyramidal neurons make en-passant synapses at subicular varicosities (Finch et al., 1983; Tamamaki and Nojyo, 1990). We show that single action-potential evoked Ca^2+^ transients were reliably activated in these varicosoties from somatic stimulation, regardless of distance from the soma (to 600 *μ*m). Using a low affinity dye that did not significantly buffer entering Ca^2+^ (Fluo-5F), Ca^2+^ transients recorded over more than 1 hour did not vary in peak amplitude or decay τ. This result implies quantal fluctuation of neurotransmission is not mediated by fluctuations in total presynaptic Ca^2+^ entry in CA1-subicular synapses, although full Ca^2+^ occupancy of syt1 is very sensitive to local Ca^2+^ placement. This result is not broadly applicable across cell types. Cerebellar granule cell varicosities show variation in total evoked Ca^2+^ entry (Brenowitz and Regehr, 2007) whereas hippocampal dentate granule cells (Jackson and Redman, 2003), cortical pyramidal cells (Koester and Sakmann, 2000) and lamprey giant axons (Photowala et al., 2005) may not.

The reliability allows calculations of buffering capacity, molar Ca^2+^ entering, and extrusion rates. Thus, if we assume a VGCC conductance of 2 pS and mean current of 0.25 pA for 1 ms during action potentials (Weber et al., 2010), then the mean number of VGCCs per varicosity is 27 with a mean channel density of 7 *μ*m^−2^. This is consistent with findings that single channels can evoke release during artificial voltage ramps (Stanley, 1993; Haydon et al., 1994; Bertram et al., 1996) and that few channels are necessary for evoked release (Bucurenciu et al., 2008; 2010). This number of channels may still allow efficient coupling of Ca^2+^ entry to exocytosis (Scimemi and Diamond, 2012). Using simulations of Ca^2+^ distribution in varicosities we investigated how this may be achieved.

We created Monte Carlo simulations (MCell) within a model varicosity to investigate Ca^2+^ entry, diffusion, and buffering. Calbindin_28K_ dominates Ca^2+^ buffering in CA1 pyramidal neuron dendrites (Müller et al., 2005) and its binding properties have been quantified with unprecedented accuracy (Nägerl et al., 2000) enabling its simulation. From experimental data, we found the transient peak [Ca^2+^]_i_ and τ were well-described by simulating calbindin_28K_ with a 2:2 ratio of high and medium-affinity Ca^2+^ binding sites. Remarkably, 3D plots of τ vs peak [Ca^2+^]_i_, vs calbindin_28K_ concentrations converge with experimental data at a calbindin_28K_ concentration identical to these neurons’ somata and dendrites (39.7 *μ*M for this study vs 40 *μ*M for somata (Müller et al., 2005)). This 2:2 ratio of binding sites (Nägerl et al., 2000) also well fit data from cerebellar purkinje neurons (Schmidt et al., 2012). Two other buffer configurations — a 3:1 ratio of sites for calbindin_28K_, and of calmodulin failed to converge with experimental data (Fig 5). Our findings, while consistent with calbindin_28k_ as a dominant buffer come with the caveat that Ca^2+^ buffering must be a function of a mix of buffers. The model also reproduced features of the experimental data, including responses to repetitive stimulation and the failure of detectable Ca^2+^ to diffuse from varicosities to their axons. Simulations were used to validate the use of exogenous dye as buffer within small varicosities. Simulating rising concentrations of the high affinity Fluo-4 dye recapitulated experimental data showing effects of buffer on measured Ca^2+^ transient decays, Ca^2+^ transient peak amplitudes, and total Ca^2+^ entering (Fig. 5).

Presynaptic VGCCs colocalize to active zones (Khanna et al., 2007) and bind SNARE complex proteins (Mochida et al., 1996; 2003; Harkins et al., 2004; Szabo et al., 2006). Furthermore, release may be activated by single VGCCs (Stanley, 1993; Bertram et al., 1996; Shahrezaei et al., 2006), although, it is unclear whether presynaptic Ca^2+^ entry occurs through channel clusters (Llinas et al., 1992; Bertram et al., 1996; Shahrezaei and Delaney, 2005) or more diffusely through a uniform distribution of channels. In the latter, individual Ca^2+^ channels might associate with primed vesicles in a 1:1 ratio. However, because Ca^2+^ signals from single VGCCs are smaller and faster than our imaging resolution, Ca^2+^ entering the terminal, even at discrete points, will appear uniformly throughout the terminal. Additionally, dye-Ca^2+^-complex diffusion may rapidly smooth signal variation. However, we have demonstrated that Fluo-5F (at 10 - 35 *μ*M) caused little perturbation of the evoked transient because most Ca^2+^ binds endogenous Ca^2+^ buffers rather than dye. (κ_dye_ = 10 to 20 for Fluo-5F vs. 75 for κ_end_). Thus, in recordings, where the terminals were close to the surface of the slice and signal-to-noise ratios were favorable, we recorded reliable localized Ca^2+^ signaling in varicosities.

These regions show faster early τ’s, requiring double exponential fits. When plotted as [Ca^2+^]_i_ vs. time, the early component τ was close to 3 ms (as fast as we can record). Peak experimental free [Ca^2+^]_i_ within these regions reached 4 *μ*M. While this does not represent concentrations responsible for exocytosis (Adler et al., 1991; Augustine et al., 1991b; Schneggenburger and Neher, 2000), the rapid decay from concentrations substantially greater than seen throughout the remainder of the varicosity indicate non-uniform VGCC distributions, consistent with channel clustering. We used MCell simulations to determine whether VGCC clustering explains stable hotspots of Ca^2+^ entry. Substantial spatial variation was only seen in simulated Fluo-5F responses if half or more of the channels in model synapses were clustered at one site, providing support to the hypothesis that channels cluster.

At CA1 pyramidal to subicular synapses, paired-pulse facilitation is apparent. This facilitation, as in other synapses (Katz and Miledi, 1968; Kamiya and Zucker, 1994; Zucker and Regehr, 2002), follows the time-course of presynaptic residual Ca^2+^. However, with measured peak evoked Ca^2+^ transients of 4 *μ*M, simulated concentrations of 10 *μ*M throughout the varicosity and local concentrations exceeding 100 *μ*M, residual free [Ca^2+^]_i_ cannot reach concentrations that will modify paired-pulse responses by a direct action on proteins that evoke synchronous release. This reiterates a number of earlier studies across many synapses (Blundon et al., 1993; Delaney and Tank, 1994; Zucker and Regehr, 2002), although Ca^2+^ signaling is necessary for short-term facilitation (Kamiya and Zucker, 1994; Mukhamedyarov et al., 2009) It has alternatively been proposed that Ca^2+^ entry following single action potentials saturates endogenous buffers (Neher, 1998) leading to enhanced subsequent transients (Jackson and Redman, 2003), or perhaps that subsequent Ca^2+^ entry is enhanced (Müller et al., 2008). In CA1 varicosities, experimentally applied paired pulse stimuli at 20 or 200 ms intervals revealed no alteration of the 2^nd^ Ca^2+^ transient, either peak amplitude or decay. Later responses in stimulus trains of 5 stimuli do show non-linear summation, which might be attributable to buffer saturation (Neher, 1998) or a secondary source of Ca^2+^ (Cochilla and Alford, 1998; Llano et al., 2000; Emptage et al., 2001; Scott and Rusakov, 2006) though notably we recorded no effect of ryanodine.

To investigate this at molecular spatio-temporal scales, simulation was used to investigate evoked Ca^2+^ transients. Throughout the varicosity, our Ca^2+^-dye buffering data and simulations demonstrate that most Ca^2+^ entering the varicosity is buffered endogenously at the timescale of imaging, similarly to results from other synapses (Koester and Sakmann, 2000; Jackson and Redman, 2003). Simulations also indicate that buffers re-release Ca^2+^ over seconds, and second stimuli force occupancy of most Ca^2+^ buffer binding sites. Supralinear rises in imaged and simulated Ca^2+^ transients after more than two stimuli indicate this can be due to buffer saturation. Simulation data indicate that total free Ca^2+^ varicosity concentrations reach 9.3 *μ*M after a single stimulus, and that buffer saturation allows a whole-terminal, paired-pulse enhancement of Ca^2+^ transients up to 1s after the first stimulus (Fig 10). However, this does not account for Ca^2+^ at scales of tens of nanometers and picoseconds in which Ca^2+^ binds to synaptotagmin1.

When Ca^2+^ transients were simulated at the nanometer scale local to syt1 molecules, it is clear that Ca^2+^ dispersal from such small regions is dominated by diffusion. Rates of removal due to buffering or local accumulation are three orders of magnitude less than this diffusion. Because diffusion dominated there was no detectable difference in the amplitudes or distribution of two transients in paired pulses at intervals of 20 or 200 ms at this scale. Indeed, modeling binding of 5 Ca^2+^ to syt1 indicates that Ca^2+^ entry within less than 50 nm of synaptotagmin causes its full occupancy. This result, along with our experimental and modeling data indicate clustering of Ca^2+^ channels may contribute to release at this synapse. Indeed such clusters of channels that are constrained to be further than 30nm from the fusion apparatus has been proposed in Calyceal synapses (Keller et al., 2015), although in those synapses Ca^2+^ requirements for release are as low as 25 *μ*M (Schneggenburger and Neher, 2000). However full Ca^2+^ occupancy of the synaptotagmin1 model is not enhanced by paired pulses. If residual Ca^2+^ is responsible for paired pulse facilitation, for which there exists a great deal of evidence (Zucker and Regehr, 2002), then these data, in addition to data indicating the prolonged binding of Ca^2+^ buffers, point to a Ca^2+^ binding site distinct from that involved in evoked release (i.e. synaptotagmin 1) as mediated by a short-term enhancement of neurotransmitter release at these synapses, such as synaptotagmin 7 (Jackson and Redman, 2003; Jackman et al., 2016). This effect is perhaps also consistent with an effect of residual bound Ca^2+^ on vesicle priming (Neher and Sakaba, 2008) and the size of the readily releasable pool (Thanawala and Regehr, 2013).

We conclude that the Ca^2+^ responsible for synchronous evoked release at en-passant pyramidal neuron varicosities reaches hundreds of micromolar concentrations at clusters of Ca^2+^ channels local to the release machinery. These clusters may represent up to half of the Ca^2+^ entry in the terminal as a whole for which the activation of fewer than 30 Ca^2+^ channels is required, but that dispersal from these sites is diffusion dominated and does not show accumulation during repetitive stimulation. Nevertheless, buffer saturation during repetitive stimulation does account for a varicosity wide enhancement of Ca^2+^ transient amplitudes which may impact short-term enhancement by recruiting other Ca^2+^ binding proteins.

## Materials and Methods

### The preparation

Experiments were performed on hippocampal slices (300 *μ*m) of male or female 20-22-day-old Sprague-Dawley rats anesthetized with isoflurane and decapitated. Hippocampi were isolated under semi-frozen Krebs Henseleit solution (in mM): 124 NaCl, 26 NaHCO3, 1.25 NaH2PO4, 3 KCl, 2 CaCl2, 1 MgCl2, 10 D-glucose, bubbled with 95% O2-5% CO2, sliced using a Vibratome. The recording chamber was superfused at 2 ml/min and maintained at 28 ± 2 °C. Experiments were performed in accordance with institutional guidelines of the University of Illinois at Chicago and the Association for Assessment and Acreditation of Laboratory Animal Care.

### Electrophysiology

CA1 pyramidal neurons were whole-cell clamped follwing visual identification using an upright microscope with an Axopatch 200A amplifier (Axon Instruments). Patch pipettes (4-5 MΩ) contained solution (in mM): potassium methane sulphonate 146, MgCl2 2, EGTA 0.025, HEPES 9.1, ATP 5 and GTP 2.5, pH adjusted to 7.2 with KOH. Pipettes were also filled with either Fluo-4 (1 mM) or Fluo-5F (200 *μ*M) and Alexa 594 hydrazide (250 *μ*M). Subicular pyramidal neurons were recorded under whole cell conditions but were held under voltage clamp to record synaptic inputs. In these latter neurons access resistance was monitored with a 10 mV voltage step before each episode. Focal stimuli (0.2 ms, 20 *μ*A or less) were applied over CA1 axons using glass-insulated monopolar tungsten microelectrodes. Cells were labeled with dye by allowing sufficient time for diffusion from the patch pipette in the live cell. Axons were tracked from the soma to their presynaptic terminals in the subiculum (Fig 1A) (Hamid et al., 2014).

### Imaging

Confocal microscopy was used to image individual varicosities of CA1 pyramidal neurons, with a 60X 1.1 NA water-immersion lens using a modified Biorad MRC 600 confocal microscope with excitation wavelengths at 488 and 568 nm (Bleckert et al., 2012). Ca^2+^-sensitive dyes of one of two different affinities to Ca^2+^ were visualized in each experiment. Dye concentration was determined by pairing these dyes with a Ca^2+^-insensitive dye (Alexa 594 hydrazide, molecular weight – 736 g·mol^−1^, identical to Fluo-5F and almost identical to that of Fluo-4 – 737 g·mol^−1^). Co-diffusion of the Fluo-4 and Alexa 594 hydrazide was demonstrated by recording absolute values of the fluorescence at axonal varicosities at rest over time. Alexa 594 hydrazide was excited with a 568 nm laser and imaged in long-pass (>580 nM). Fluo dyes were separately excited with a 488 nm laser and imaged in bandpass (510-560 nm). Images were taken separately to ensure no cross channel bleed-through.

Fluo-4 and Alexa 594 hydrazide signals at varicosities were identified within 20 mins of whole-cell access in 10 neurons. Signal strength was normalized to this time point for both dyes. A comparison of the increase in dye intensity over the following 20 mins with no stimulation reveals strong correlation with a slope of 1.01 (Fig 10B, this fit was forced through the origin because both dyes must be at a concentration of zero at the start of the experiment. Without this, the slope was 0.94). As an independent control to confirm that the slight difference in molecular weights alone does not alter rates of diffusion of the two dyes in the axon an MCell model was constructed of axons (diameter, 0.12 *μ*m, varicosities at 4 *μ*m intervals). Release of dyes were simulated within a 10 *μ*m diameter soma and diffusion to varicosities up to 500 *μ*m from the soma were simulated. There was no significant difference between rates of diffusion of the two dyes and diffusion times to varicosities were within times seen for experimental data (supplementary Fig 1) in which dye diffusion into varicosities was monitored using the Alexa 594 hydrazide fluorescence signal. Thus, both dyes reach the varicosity at the same rate, which allows the use of Alexa 594 hydrazide as a “standard candle” for measuring dye concentration. Thus, Alexa 594 hydrazide fluorescence in axon varicosities was used to determine the concentration of Ca^2+^-sensitive dye in the terminal by calculation of that fluorescence as a fraction of its fluorescence in the recording pipette where its concentration was known.

The fluorescence intensity of Alexa 594 hydrazide fluorescence was measured throughout the experiment in the terminal using fixed parameters on the imaging system. A plot of intensity against time approached an asymptote towards 60 mins after obtaining whole-cell access. The absolute fluorescence at the electrode tip was compared to that of the axon varicosities. At the end of the experiment the axon typically blebbed to ∼5 *μ*m—large enough for the microscope point spread function to allow for absolute fluorescence of the axon to be measured. This phenomenon was never present during the recording of stimulation-evoked Ca^2+^ transients. Its occurrence was observed subsequent to a rise in resting Ca^2+^ seen after about an hour of recording. Fluorescence in the tip of the pipette where the dye concentration was known was measured in the tissue at the same depth as the axon. This allowed calculation of the axon dye concentration after the experiment ended. It was then straightforward to compare all previously measured values of Alexa 594 fluorescence, to give absolute dye concentrations throughout the experiment. Recordings in which all of these criteria could not be met were rejected from analysis. Absolute Ca^2+^ concentrations were calculated in each varicosity using equations 1 and 2 (below). For these calculations we obtained saturated Fluo-4 or Fluo-5F intensity values in varicosities at the end of the experiment by repetitive stimulation and calculated minimum fluorescence values determined from the data in Figure 1C.

Ca^2+^ binding properties of Fluo-4 and Fluo-5F were determined using Ca^2+^ standards (Invitrogen) at 28 ± 2 °C (the temperature at which experiments were performed) and a pH of 7.2 (to which the intracellular patch solutions were buffered). Log plots of these data points were used to determine K_d_ (Fig 1C). Calibrated measurements of dye fluorescence within neurons were also made for each dye (n=2 for Fluo-5F and n=2 for Fluo-4). Whole-cell recordings were obtained in which the patch electrode contained either Fluo-4 or Fluo-5F. Whole cell access was maintained until soma and dendrites were clearly labeled. The electrode was then carefully withdrawn. Baseline fluorescence intensities were measured. The Ca^2+^ ionophore, ionomycin (5 *μ*M) was added to the superfusate. Fluorescence intensities were measured at 2 min intervals until the signal reached a stable maximum. The superfusate was then replaced with a solution containing 0 Ca^2+^ and 10 mM EGTA and images taken until the fluorescence intensity reached a stable minimum. This solution was then replaced with a solution containing buffered Ca^2+^-EGTA ([Ca^2+^] = 0.78 *μ*M) and fluorescence in the neuron was measured. Ratios of maxima to minima were very close to those obtained with standards. The standard data points were then plotted over the log plots obtained *in vitro*. A small correction was applied to the calculated Kd for Fluo-4 (black line, Fig 1B) but the data obtained with Fluo-5F gave a value of Kd that was not measurably different than that obtained *in vitro*. These values (Fluo-4 Kd= 0.44 *μ*M, Fmin/Fmax = 0.066 and Fluo-5F Kd = 1.49 *μ*M, F min/F max = 0. 023) were used in all subsequent calculations.

### Effects of introduction of buffers to cell compartments

It is possible to calculate the intracellular Ca^2+^ concentration ([Ca^2+^]_i_). For a non-ratiometric dye with a Hill coefficient of 1, [Ca^2+^]_i_ is determined from equation 1:

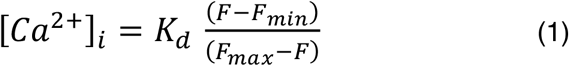

The Ca^2+^ dye minimum fluoresence intensity (F_min_) was calculated as a ratio of F_max_ determined from the dye calibration results summarized in figure 1B, from each cell at the end of the experiment. Absolute values of F_min_ and F_max_ were corrected by the observed value of Alexa 594 for each time point as a ratio of its value at the end of the experiment when F_max_ was measured. A corrected value of Ca^2+^ dye fluorescence in the varicosity (F) was calculated from the measured varicosity fluorescence (Fmeas) at each time point used for analysis and then re-expressed as a ratio of F_max_, corrected by comparison to the Alexa 594 signal throughout the experiments. This is because F_max_ was determined at the end of the experiment and consequently needed to be scaled for each time point at which measurements were made during the experiment. Thus, F is given by:

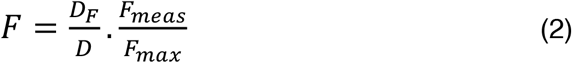

where *D* is the Alexa 594 fluorescence at each time, and *D_F_* is the final Alexa 594 fluorescence. Thus, for experiments using Fluo-4 or Fluo-5F, where F and F_min_ in equation (1) are ratios of F_max_, and F_max_ = 1, [Ca^2+^]_i_ was calculated as follows:

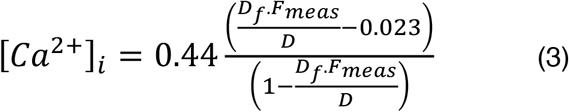

These experiments required constant laser intensity and recording parameters throughout the experiment. To minimise photobleaching, imaging was performed only transiently during evoked responses (1s per stimulus, < 15 s total per experiment). To use calcium-sensitive dyes as buffers to investigate the fate of Ca^2+^ that enters presynaptic terminals on stimulation, we must calculate their buffering capacities (k_dye_) in the cytosol of the terminal. Since each molecule of dye binds just one Ca^2+^ ion, the Hill equation with a coefficient of one can be used to calculate κ_dye_ over a change in Ca^2+^ concentration (Δ[Ca^2+^]_i_) from [Ca^2+^]1 to [Ca^2+^]_2_.

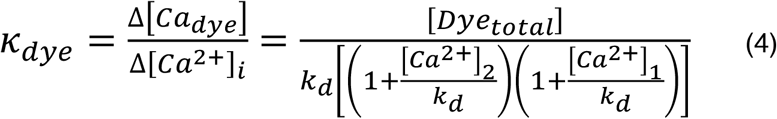

where [*Ca_Dye_*] is the concentration of Ca^2+^-bound dye, and [*Dye_total_*] is the total dye concentration. Note that this approach takes into account the change in [Ca^2+^]_i_ in the pyramidal cell terminals during the stimulus which is large (approx. 1 *μ*M). Other approaches with smaller Ca^2+^ changes use resting [Ca^2+^]_i_ as a basis for calculating κ_dye_ (Neher and Augustine, 1992).

When a rapid Ca^2+^ pulse enters a cell compartment, free Ca^2+^ may be removed first by binding to intracellular endogenous buffers, and possibly by diffusion into neighboring compartments, and then by pumps. We have used methods used by Jackson and Redman (Jackson and Redman, 2003) originally described by Melzer et al (Melzer et al., 1986) to determine the buffering characteristics of Ca^2+^ in CA1 pyramidal neuron presyaptic varicosities. From these methods we can determine the quantity of calcium entering the varicosity, the mean free [Ca^2+^]¡ within the varicosity immediately after the stimulus, the endogenous Ca^2+^ buffering capacity, and the rate of removal of Ca^2+^ from the cytosol. From this we have developed simulations of Ca^2+^ entry, diffusion, and buffering in the presynaptic terminal.

The relationship between Ca^2+^ unbinding rates from the dye and rebinding either to dye or endogenous buffers can be used to calculate endogenous buffering capacity (κ_end_) of the terminal. If the value of κ_dye_ varies during the experiment then we assume a constant rate of Ca^2+^ extrusion from the terminal (τ_ext_). The decay rate (τ) of a pulse-like Ca^2+^ signal in a cell compartment is described by the equation:

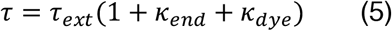

Thus, we obtained values of κ_end_ by fitting equation (5) to plots of τ from experimental data vs κ_dye_ from equation (4). This approach does have drawbacks, if processes modifying Ca^2+^ removal or adding to cytosolic Ca^2+^ occur after action potentials. Such processes include diffusion of the Ca^2+^-dye complex from the measured compartment, or release of Ca^2+^ from internal stores. In equation (5) these effects are grouped into a single variable k_end_. Nevertheless, the result (κ_end_) can be obtained independently of computed absolute values of Ca^2+^, or even of background fluorescence meaurement errors. It therefore serves as an independent measure of whether the following measurements of k_end_ are reasonable.

By measuring the peak amplitude of the free Ca^2+^ transient throughout the varicosity over a range of values of κ_dye_, we may assume that the total change in Ca^2+^ concentration due to a stimulus is described by the equation:

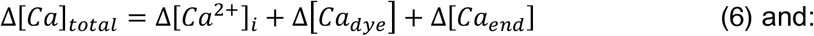

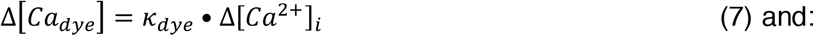

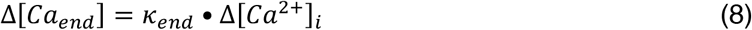

Where [*Ca_end_*] is the concentration of endogenous buffer bound to Ca^2+^ and *Δ[Ca]_total_*is the total stimulus-evoked change in calcium concentration in the cell compartment. Thus combining these equations we may state:

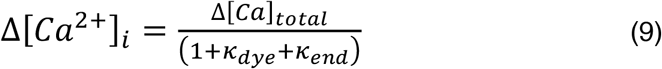

Where Δ[Ca^2+^]_i_ varies with k_dye_. Values of Δ[Ca^2+^]_i_ are computed from our data using equation (3) and values of κ_dye_ from equation (4). For each action potential, as the value of κ_dye_ rises it will come to dominate binding of Ca^2+^ entering the cell compartment. This approach is useful because the value of Δ[Ca_dye_] can be calculated for each action potential in each presynaptic terminal as the value of κ_dye_ increases by diffusion of dye from the soma. Equations (6), (7), and (8) can also give:

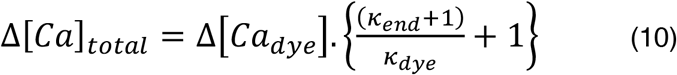

Values of Δ[Ca]_total_, and κ_end_ can be determined by extracting constants from fits of either equation (9) or (10). The true value of peak Δ[Ca^2+^]_i_ in the varicosity when no dye is present is obtained by extrapolating the fit to the y intercept in equation (9) where κ_dye_ = 0. To calculate total Ca^2+^ entering the terminal from the concentrations obtained from either equation (9) or (10), we calculated varicosity volumes from images using Alexa 594 hydrazide. Varicosities are larger than the smallest structures that can be imaged in our microscope. Point spread data from 0.2 *μ*m latex microspheres were determined by imaging under the same light path as all data in this study (568 nm excitation, long pass emission). The point spread was Gaussian in x-y and z dimensions with an x-y dimension half maximal width of 0.45 *μ*m.

Varicosities approximated elipsoids with the long axis along the line of the axon. We measured length (l) and width (w), assuming depth was similar to the width because it was not possible to obtain sufficient z-plane resolution to accurately determine depth. Mean measured varicosity length (l) was 2.3 ± 0.2 *μ*m and width (w) was 1.2 ± 0.1 *μ*m. These values are quite similar to terminal sizes obtained from electron microscopic images (Harris and Weinberg, 2012).

Assuming the varicosities were ellipsoid, volume of the varicosity is given by:

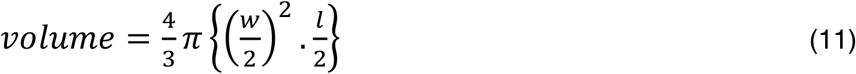

Chemicals were obtained as follows: Alexa dyes, Fluo-4, Fluo-5F, Ca^2+^ standards; (Thermo Fisher, Eugene OR), salts, buffers etc; Sigma (St Louis MO). Fits to datasets were performed in Igor Pro (Wavemetrics, IA USA). Errors bars from fitted data represent the 95% confidence limits of those fits. Otherwise errors are reported as the standard error of the mean. Significance was tested with Student’s t test or 2 factor ANOVAs where appropriate.

### Simulations

Monte Carlo simulations were applied to Ca^2+^ buffering within a model of the CA1 axon varicosity based on the data obtained experimentally in this study. Simulations were run in the MCell environment (Kerr et al., 2008) in which a 3D mesh model of the presynaptic terminal was created based on measurements determined from these experiments and from electron microscopic images of hippocampal presynaptic varicosities (Harris and Weinberg, 2012). Ca^2+^ entry, diffusion binding and removal from the terminal were modeled using initial parameters obtained from experimental data in this study, and from the literature and from published sources. Possible Ca^2+^ binding proteins and their concentrations were investigated by comparing multiple parameters from experimental data and the results of simulations. All of the parameters used are outlined in tables 1, 2, and 3. Animated visualizations of these simulations are shown in Supplementary materials.

## Acknowledgements

We would like to thank Drs Janet Richmond and Nelson Spruston for helpful discussions throughout the course of this study. This work was funded by NIH grants R01 MH084874 and R01 NS052699 to SA and F31 NS063662 to EH.

